# Spatial transcriptional signatures define margin morphogenesis along the proximal-distal and medio-lateral axes in the complex leaf of tomato (*Solanum lycopersicum*)

**DOI:** 10.1101/772228

**Authors:** Ciera C. Martinez, Siyu Li, Margaret R. Woodhouse, Keiko Sugimoto, Neelima R. Sinha

**Affiliations:** Department of Molecular and Cellular Biology, University of California at Berkeley, Berkeley, CA 94709, USA; Berkeley Institute of Data Science, University of California at Berkeley, Berkeley, CA 94709, USA; Department of Plant Biology, University of California at Davis, Davis, CA 95616, USA; RIKEN Center for Sustainable Resource Science, Tsurumi, Yokohama, 15 230-0045 Japan

## Abstract

Leaf morphogenesis involves cell division, expansion, and differentiation in the developing leaf, cells at different positions along the medio-lateral and proximal-distal leaf axes divide, expand, and differentiate at different rates. The gene expression changes that control cell fate along these axes remain elusive due to difficulties in precisely isolating tissues. Here, we combined rigorous early leaf characterization, laser capture microdissection, and transcriptomic sequencing to ask how gene expression patterns regulate early leaf morphogenesis in wild-type tomato (*Solanum lycopersicum*) and the leaf morphogenesis mutant *trifoliate*. We observed transcriptional regulation of cell differentiation along the proximal-distal axis and identified molecular signatures delineating the classically defined marginal meristem/blastozone region during early leaf development. We describe the importance of endoreduplication during leaf development, when and where leaf cells first achieve photosynthetic competency, and the regulation of auxin transport and signaling along the leaf axes. Knockout mutants of *BLADE-ON-PETIOLE2* exhibited ectopic shoot apical meristem formation on leaves, highlighting the role of this gene in regulating margin tissue identity. We mapped gene expression signatures in specific leaf domains and evaluated the role of each domain in conferring indeterminacy and permitting blade outgrowth. Finally, we generated a global gene expression atlas of the early developing compound leaf.

**One-sentence summary:** Rigorous structural characterization, laser capture microdissection, and transcriptomic sequencing reveal how gene expression patterns regulate early morphogenesis of the compound tomato leaf.

The author responsible for distribution of materials integral to the findings presented in this article in accordance with the policy described in the Instructions for Authors (www.plantcell.org) is Ciera C. Martinez.

## INTRODUCTION

A major theme in plant development is the reiteration of patterning events, which are influenced by the identity and relative arrangement of neighboring plant parts. The phytomer concept describes reiterated units of the leaf, stem, and axillary bud that make up the aboveground shoot (Sussex and Kerk, 2001). Molecular analyses comparing development in various plant species suggest that the reiteration of developmental patterning in plants is defined by the recruitment of a common molecular toolbox and the dizzying array of leaf architecture found in plants results from variations on a common genetic regulatory program (Tsukaya, 2014; Bendahmane and Theres, 2011; Blein et al., 2008).

The shoot apical meristem (SAM), which is located at the growing tip of the shoot, is a dome-like structure containing reservoirs of continually self-renewing stem cells and is characterized by spatially defined zones. The peripheral zone of the SAM gives rise to most lateral organs, including leaves. Like the SAM, the angiosperm leaf has been historically defined in terms of zones and spatial cell organization. Leaf development begins with periclinal cell divisions on the periphery of the SAM and continues as cells proceed through the specific steps of development beginning with cell division, followed by cell expansion and cell specialization. In many instances, this specialization involves endoreduplication. The timing of these stages varies depending on the cell position on the leaf primordium.

Leaf morphogenesis and patterning occur along three main axes: the abaxial-adaxial, proximal-distal, and medio-lateral axes. Many studies have focused on the importance of the abaxial-adaxial boundary in establishing leaf polarity (Eshed et al., 2001; Moon and Hake, 2011; Kidner and Timmermans, 2007), but less is known about the proximal-distal and medio-lateral axes of the leaf. During the development of most eudicot leaves, cells differentiate more rapidly in the distal (top) region than in the proximal (base) region. Along the medio-lateral axis, the differentiation at the margin of a leaf is decelerated relative to the more medial regions (midvein, rachis, petiole). Thus, historically, the leaf margin is of particular interest because it maintains cellular pluripotency longer than the other regions and has even been described as a meristematic region termed the marginal meristem (Poethig and Sussex, 1985b; Avery, 1933) or marginal blastozone (Hagemann and Gleissberg, 1996).

Although the developmental fate, homology, and even the name of the margin region of a leaf have been debated for roughly 100 years, there is general agreement that the process of cell differentiation in this region largely determines final leaf shape (Ori et al., 2007; Efroni et al., 2008; Scarpella and Helariutta, 2010). The regulation and modulation of the margin identity of a leaf are responsible for blade expansion, serrations, lobing, vascular patterning, and new organ initiation, as in the case of leaflet initiation in compound leaves (Scarpella et al., 2010; Bilsborough et al., 2011).

The genetic regulation and coordination of leaf morphogenesis involve distinct changes in gene expression, as revealed by leaf transcriptomic studies in spatially defined regions across the proximal-distal axes of the simple-leaved plant *Arabidopsis thaliana* (*A. thaliana*) (Beemster et al., 2005; Andriankaja et al., 2012; Efroni et al., 2008). These studies revealed the role of endoreduplication (DNA replication without cell division) in the acquisition of leaf morphogenic potential (Beemster et al., 2005; Andriankaja et al., 2012; Efroni et al., 2008). The transcriptional mapping of gene expression changes in *A. thaliana* (Beemster et al., 2005; Efroni et al., 2008; Andriankaja et al., 2012), *Solanum lycopersicum* (tomato) (Ichihashi et al., 2014), and *Zea mays* (maize) (Li et al., 2010) shed light on how patterning by cellular differentiation along the proximal distal axis is established. However, this information has not yet been precisely mapped at the transcriptome level with sufficient spatial resolution to define margin and midvein/rachis/petiole transcriptional identity.

Interestingly, the tomato mutant *trifoliate* (*tf-2*) loses morphogenetic competence during early leaf development and only produces three leaflets: a terminal leaflet and two lateral leaflets subtended by a long petiole (Robinson and Rick, 1954; Naz et al., 2013). The *tf-2* phenotype is caused by a nucleotide deletion resulting in a frameshift in the translated amino acid sequence of an R2R3 MYB transcription factor gene (Solyc05g007870) (Naz et al., 2013). Histological and scanning electron microscopy (SEM) analyses of the *tf-2* mutant revealed that the marginal blastozone region is narrower and has fewer cells, a three-fold increase in epidermal cell size, and faster cell differentiation than the wild type (Naz et al., 2013). While the application of auxin to the margins of wild-type *S. lycopersicum* leaf primordia causes leaflet initiation (Koenig et al., 2009; Naz et al., 2013), in *tf-2*, the margin is unable to generate leaflets in response to exogenous auxin, indicating that it lacks organogenic competency during early development (Naz et al., 2013). Understanding why this mutant is incapable of initiating more than two lateral leaflets, while wild-type leaves continue to generate an average of ten leaflets at maturity (Naz et al., 2013), could help reveal the mechanisms regulating margin maintenance and identity during complex leaf development.

Here, we used the complex tomato leaf as a model system to study the transcriptional mechanisms directing spatial cell differentiation processes during a key developmental stage in a young leaf, including the establishment of margin identity, proximal-distal patterning, and leaflet initiation. Since a leaf primordium develops at varying rates in a spatially defined manner, different developmental stages can be observed at the same time in a single leaf (Hagemann and Gleissberg, 1996; Ori et al., 2007). We anatomically characterized the earliest developmental stages in tomato and identified leaf age P4 as the stage at which the medio-lateral and proximal-distal axes are first identifiable while also containing multiple stages of leaflet organogenesis. We also characterized the role of endoreduplication in tomato leaf morphogenesis. To map the spatial transcriptional regulation of the P4 leaf using Laser Capture Microdissection (LCM), we isolated six highly specific tissues previously unattainable during early tomato leaf development and performed RNA-seq analysis to identify gene expression changes that accompany the establishment of spatial cell differentiation patterning during leaf organogenesis. We also included *tf-2* in our analysis, as *tf-2* lines have early loss of morphogenetic potential in the leaf margin, thus helping us uncover a cluster of genes whose expression differs only in regions that define organogenetic capacity in the margin at the P4 stage. We further validated our results through molecular visualization, providing the first evidence for when and where photosynthesis begins in a leaf. We also used CRISPR knockout lines to explore the role of BLADE-ON-PETIOLE2 (BOP2) (Solyc10g079460) in margin development. Our approach allowed us to predict multiple gene expression differences that help explain the molecular identity of the classical described, but never transcriptionally defined, marginal meristem/blastozone region and built the first global transcriptome atlas of an early developing compound leaf.

## RESULTS

### Characterization of the P4 stage of tomato leaf development

The goal of this work was to characterize gene expression changes that occur during tomato leaf morphogenesis. To define the scope of this our work, we focused on the medio-lateral axis in an attempt to identify how the marginal blastozone maintains the potential for leaflet organogenesis and the regulation of cell fate identity. We chose to use the leaf stage primordium 4 (P4), the fourth oldest leaf emerging from the apical meristem (**Figure 1A and B**). P4 provides a comprehensive snapshot of tomato leaflet development, as it contains three distinct stages of leaflet development. During this stage, the most distal region (destined to become the terminal leaflet) is undergoing early blade expansion, while the most proximal region is undergoing lateral leaflet initiation, and central to these positions is the recently initiated lateral leaflets. All three regions can be anatomically defined, allowing the boundaries along both the medio-lateral and proximal-distal axes to be clearly delineated. With our scope defined, we began our analysis with a systematic survey of tissue differentiation patterns of the P4 leaf using a combination of scanning electron microscopy (SEM) and histological approaches to establish the cellular context for detailed tissue-specific gene expression analysis.

**Figure 1.**
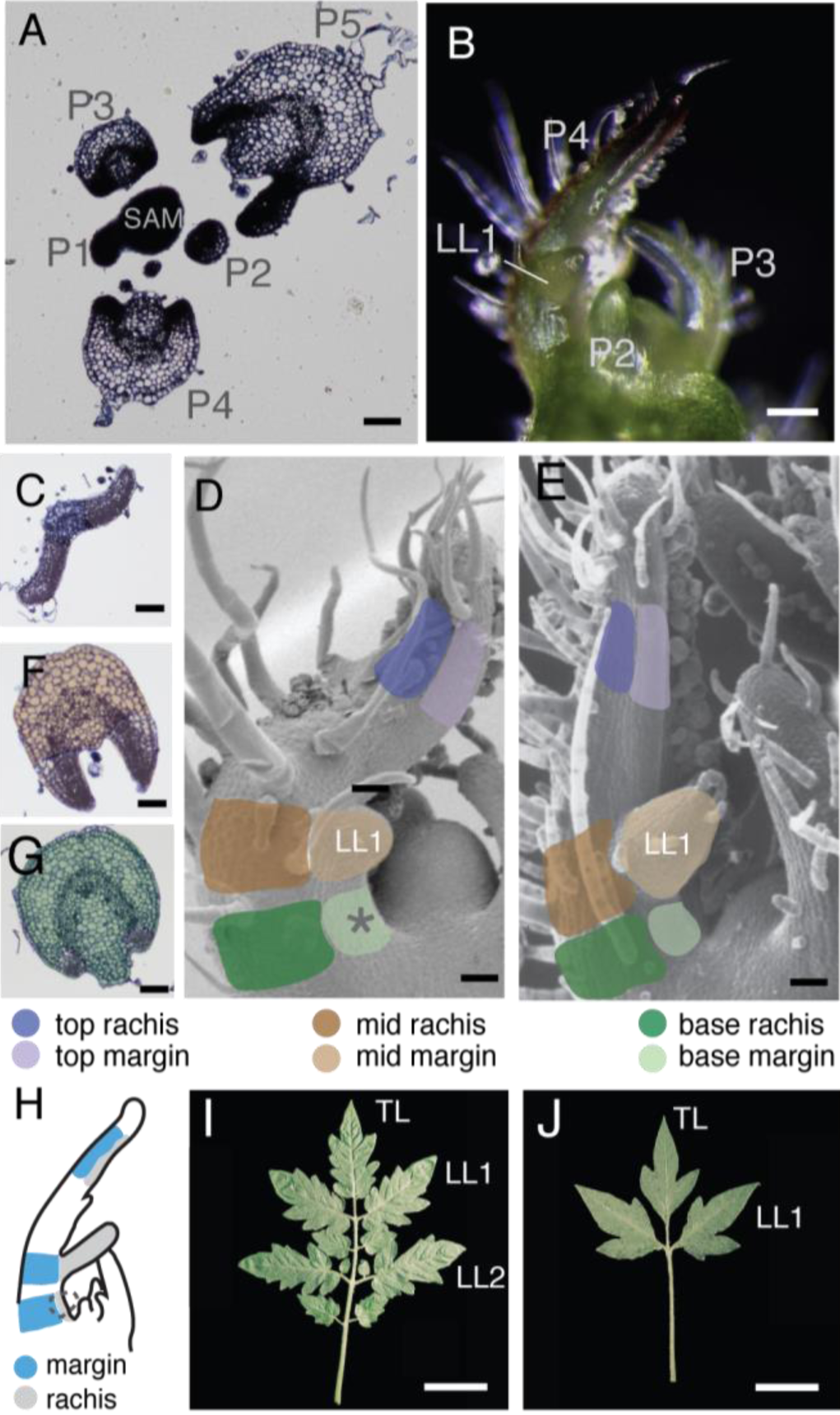
Experimental set-up for sampling *S. lycopersicum* P4 leaves. (A) Transverse section from a wild-type apex showing leaf primordia P1-P5 in relation to the Shoot Apical Meristem (SAM). (B) Image of a wild-type apex during P4 leaf development. Images of transverse sections from the (C) top, (F) middle, and (G) base regions of a wild-type P4 leaf. Colors highlight the separation of the margin (lighter colors) and rachis (darker colors) along the top (purple), middle (brown), and base (green). Schematic diagram of a P4 leaf illustrating the six regions identified in (D) wild type and (E) *tf-2*. (H) Schematic diagram showing how the margin (gray) and rachis (blue) of a leaf were defined in this study. Images of leaves from wild type (I) and *tf-2* (J). Scale bars (A-E) = 100 μm and (I) and (J) = 5 mm.

We defined three distinct regions of the P4 leaf along the proximal-distal axis, which are referred to as the top, middle, and base hereafter (**Figure 1C-G**). These three regions can further be divided into two distinct tissues types that define the medio-lateral axis: the margin and the midrib/midvein/rachis, hereafter termed the rachis for brevity (**Figure 1C,G, and F**). The most distal region, the top, will ultimately become the terminal leaflet of the mature leaf (**Figure 1C**). In P4 leaves, the top margin region has already begun to develop lamina tissue (blade) and has not yet developed any tertiary vasculature, but the future midvein in the top contains vascular cells including xylem and phloem (**Figure 1C**). The middle margin tissue has initiated the first lateral leaflets (henceforth called LL1), the first leaflets to form from the marginal blastozone, and the rachis tissue displays clear vascular bundles and more than four layers of cortex cells (**Figure 1F**). The most proximal area is the base, where rachis tissue has established vascular bundles (**Figure 1G**). Cells in the margins of all three regions along the proximal-distal axis are small and non-vacuolated and have likely undergone little elongation, a characteristic of marginal blastozone tissue (Hagemann and Gleissberg, 1996) (**Figure 1C, F, and G**). Tomato leaflets initiate in pairs proximal to previous leaflet initiation sites, and therefore, the next leaflets to arise, Lateral Leaflets 2 (LL2), will occur at the base margin region of a P4 leaf (**Figure 1D**).

To further delineate margin identity, we also characterized *tf-2*, a tomato mutant unable to initiate leaflets past LL1, to compare its margin identity and marginal organogenesis capacity with the wild type (**Figure 1E and J**). The developmental fate of the *tf-2* mutant diverges from that of the wild type at P4, as the margin is unable to form leaflets after LL1. Therefore, comparing *tf-2* and wild type allowed us to explore two leaves of comparable developmental age but with different organogenic potential, i.e., different abilities to form leaflets. The anatomical characterization of wild type and *tf-2* revealed precise cell types present across a P4 leaf, serving as a proxy for defining cell differentiation.

### Cell division and endoreduplication in the P4 leaf

Previous transcriptomic studies tracing proximal-distal cell division patterning and cellular processing indicated that changes in gene expression are responsible for the regulation of cell division, cell elongation, and endoreduplication during differentiation in developing *A. thaliana* leaves (Beemster et al., 2005; Efroni et al., 2008; Andriankaja et al., 2012; Donnelly et al., 1999). Endoreduplication is thought to be a defining component of *A. thaliana* leaf morphogenesis (Beemster et al., 2005; Gutierrez, 2005), with ploidy levels varying from 2C to 32C (Melaragno et al., 1993; Beemster et al., 2005). Endoreduplication occurs at the onset of leaf differentiation, and elongation occurs after cell proliferation, when cell ploidy levels increase due to successive rounds of DNA replication, often resulting in increased cell size (Kondorosi et al., 2000; Sugimoto-Shirasu and Roberts, 2003; De Veylder et al., 2011). While endoreduplication occurs at rates (256C to 512C) during tomato fruit development (Bergervoet et al., 1996; Joubès et al., 2000; Cheniclet et al., 2005; Bourdon et al., 2010), where and to what extent endoreduplication occurs during tomato leaf development are currently unknown. Since no reports were available on endoreduplication during tomato leaf development, we carefully characterized cell division and endoreduplication processes at the P4 stage to identify similarities and differences between early leaf development in tomato and *Arabidopsis*.

To observe where cell division is occurring throughout the P4 leaf, we used 5-ethynyl-29-deoxy-uridine (EdU), which is incorporated during the S phase of the cell cycle and serves as a proxy to map cell division locations. Along the mediolateral axis, in wild type and to a lesser extent in *tf-2*, EdU fluorescence was more prominent in the margin compared to rachis tissue (**Figure 2E-F**), indicating that the margin tissue is actively undergoing cell division, as expected for marginal blastozone tissue. At the base margin region of wild type, where Lateral Leaflet 2 (LL2) will arise, EdU was incorporated in a cluster (**Figure 2E**), clearly demonstrating early cell division processes during LL2 initiation. Therefore, during early P4 development, LL2 initiation has already begun, although this is not always obvious based on external views of the leaf (**Figure 1B and D**). The *tf-2* mutant did not show clustering of EdU fluorescence in the base margin (**Figure 2E and F**), revealing that the cell divisions needed for LL2 initiation have not occurred. In conclusion, cell division across the mediolateral axis in wild type and *tf-2* reflects similar processes that occur in *A. thaliana* (Donnelly et al., 1999), *where cells are actively dividing in the margin. The cell divisions needed for LL2 initiation at P4 have already begun in the wild type but not in tf-2*. Therefore, the mechanism that restricts LL2 initiation in *tf-2* is likely in place at the P4 stage of development.

**Figure 2.**
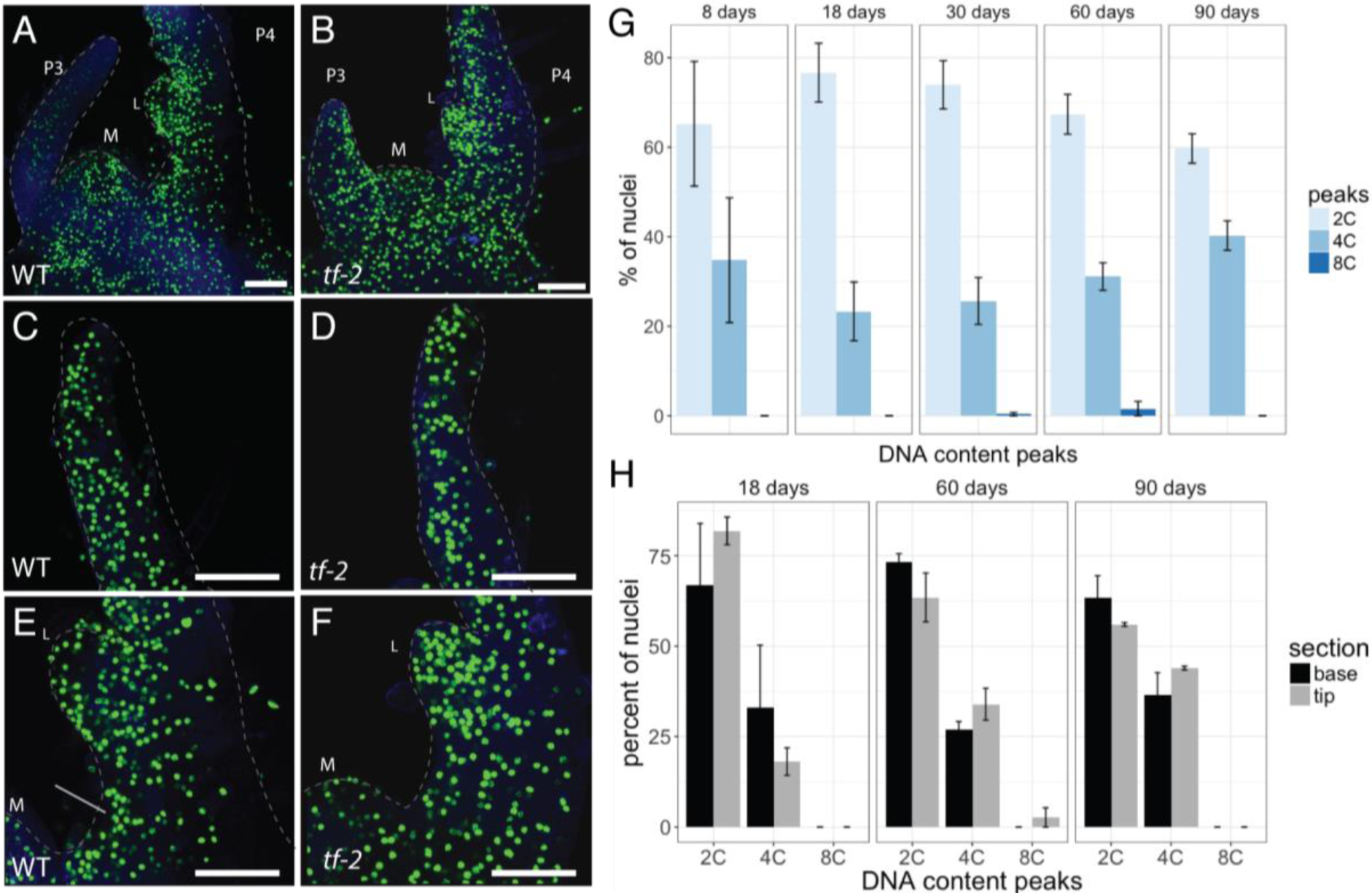
Characterization of the cell cycle using EdU fluorescence and flow cytometry. (A-F) Representative confocal images using EdU fluorescence to detect sites of cell division at the shoot apex of (A, C, E) wild type and (B, D, F) *tf-2* plants. Images in (C-D) show the top region of a P4 leaf, while (E-F) show the middle and base regions of a P4 leaf. Arrow in (E) points to clustering of EdU fluorescence suggesting sites of active cell division needed for leaflet initiation. (G) Bar graph showing DNA content peaks based on flow cytometry of leaf tissue. Tissue was collected from the oldest leaf of the plant at 8-90 days after germination. (H) Bar graph displaying the DNA content peaks from flow cytometry comparing leaf tissue sampled from the base (black) and tip (gray). Error bars show the standard deviation across at least three replicates at each time point. Scale bars = 100 μm.

We used flow cytometry to measure DNA contents in tissues from the terminal leaflets of leaves across several developmental ages. We grew all plants at Day 0 and sampled the oldest leaf on the plant at each timepoint. Due to the limited availability of tissue from the youngest leaf, we performed flow cytometry of whole terminal leaflet tissue beginning at 8 days old (P6 stage leaf). We detected a combination of 2C and 4C nuclei at all stages examined (**Figure 2G**). The 4C nuclei were likely G2 nuclei observed following DNA replication and did not reflect the endoreduplication process, although a few 8C nuclei were present at 30 and 60 days, perhaps representing cell-type-specific endocycling (**Figure 2G**). In *A. thaliana* plants, there is a difference in ploidy level between tip and base cells (Skirycz et al., 2011), but this was not the case in our analysis (**Figure 2H**). We conclude that endoreduplication is not as pronounced in tomato as in *Arabidopsis* and is likely not a vital aspect of tomato leaf morphogenesis, illustrating the diversity of cellular processes in leaf morphogenetic strategies between species.

### Laser capture of six regions of the P4 tomato leaf

Since the P4 leaf is representative of two key developmental processes that define leaf development, i.e., margin vs. rachis specification and leaflet initiation and morphogenesis, we analyzed the P4 stage more carefully. We took advantage of our comprehensive anatomical characterizations to generate a map delineating the medio-lateral axis and leaflet organogenesis. We employed Laser Capture Microdissection (LCM) following explicit rules for tissue collection (**S1 Figure and Movie 1**) on P4 leaves of both wild type and *tf-2* lines to capture gene expression differences that might explain the morphogenetic differences in the margins of *tf-2* plants. Specifically, we sectioned tomato apices transversely to isolate the same six sub-regions in both wild type and *tf-2*, including the (1) top margin blastozone region (top margin), (2) top rachis, (3) middle margin, (4) middle rachis, (5) base margin, and (6) base rachis (**Figure 1C-G, Movie 1**). We attempted to collect enough tissue for seven replicates per sample, but due to the fragility of RNA at such a small tissue size, a few replicates did not pass quality control and were lost during various steps in the pipeline, resulting in a total of 3–6 biological replicates per region. We collected tissue from 6–8 apices per biological replicate to obtain a minimum of 2 ng of RNA per replicate. The number of cuts needed to achieve minimum RNA levels varied depending on sample and tissue density, and the total tissue area collected also varied among samples (**S2 Figure B and D**). The isolated mRNA from the collected tissues was further amplified and prepared for Illumina sequencing (see Methods). Each replicate resulted in an average of 4.9 million sequencing reads (**S2 Figure A and C**). To assess the overall similarity between samples, we visualized gene expression using Multidimensional Scaling (MDS) for each of the six subregions per genotype. Tissue types in each genotype generally clustered together in multidimensional space (**S2 Figure A**) and we observed an additional separation of margin and rachis tissue regions (**S2 Figure C**).

### Differential gene expression between wild-type margin and rachis tissue along the proximal-distal axis reveals signatures of morphogenetic states during early leaf development

Identifying genes that are differentially regulated in margin versus rachis tissue in each region would shed light on gene expression patterning along the medio-lateral axis. To explore the differences between margin and rachis tissue in the three regions along the proximal-distal axis, we performed pairwise differential gene expression analysis of wild-type samples comparing the margin and rachis in each region (top, middle, base) separately (**Dataset S1**) using edgeR (Robinson et al., 2010). We then performed Gene Ontology (GO) enrichment analysis of the significantly up-regulated genes in these samples (BH-adjusted p value < 0.05) (**Dataset S2**). More upregulated genes in the margin region, which has historically been considered to proceed at a slower rate through the morphogenetic stages, were enriched in GO terms associated with cell processes that occur during early morphogenesis compared to rachis tissue. For example, when we looked for differentially expressed genes between the margin and rachis tissue in the top region, we identified 603 genes that were up-regulated in the margin (**Figure S3 A**). These genes were enriched in GO terms that likely reflect cell division processes, including chromatin and DNA processing (**Figure 3A, Dataset S2**). Conversely, up-regulated genes in the rachis were enriched in GO terms reflecting the cell specialization stage of morphogenesis, including transport, photosynthesis, sugar biosynthesis, and carbohydrate metabolism (**Figure 3A-C, Dataset S2**).

**Figure 3.**
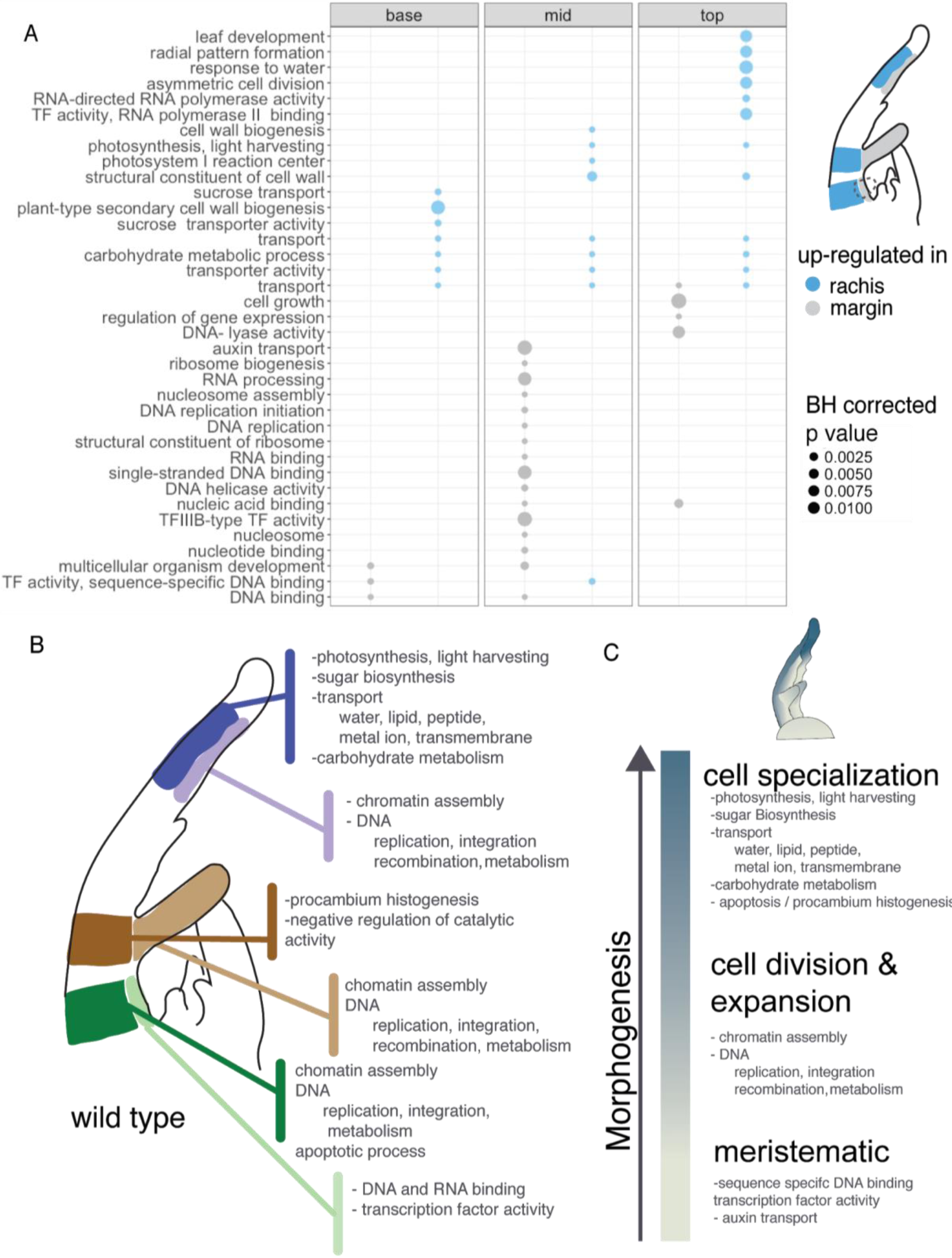
Pairwise differential gene expression between the rachis and margin in each region along the proximal-distal axis in a wild-type P4 leaf. (A) Graph summarizing representative enriched GO terms describing the significantly up-regulated genes in each region (top, mid, and base) from differential gene expression analyses performed on wild-type plants. Point size represents Benjamini-Hochberg (BH) corrected P-values in margin (gray) and rachis (blue) tissue. (B) Schematic diagram summarizing enriched GO terms of differentially up-regulated genes in each region of the P4 leaf. Colors highlight the separation of the margin (lighter colors) and rachis (darker colors) along the top (purple), middle (brown), and base (green). (C) Schematic diagram showing GO categories that help define each morphogenetic state along the leaf.

A comparison of genes expressed in the margin and rachis in the most proximal region (the base) revealed 1722 genes that were up-regulated in the rachis, which were enriched for GO terms related to cell division, chromatin assembly, and DNA processing. By contrast, only 94 differentially expressed genes were down-regulated genes in the margin region vs. rachis at the base (**S3 Figure A**). These downregulated genes were enriched for GO terms related to transcription factor activity and auxin influx (**Figure 3A, Dataset S2**). The types of genes that were differentially expressed between the margin and rachis also appeared to reflect that stage of morphogenesis of each region and perhaps the distal-to-proximal wave of differentiation (**Figure 3B and C**). The up-regulated genes in the top and middle margin regions were enriched in GO terms describing active RNA, DNA, and chromatin processing, whereas those in the base were enriched in GO terms similar to those in the rachis. The active processing of RNA, DNA, and chromatin are key gene expression signatures of cell division and expansion. The base region of the P4 leaf is still in these middle stages of morphogenesis and just beginning to start secondary cell wall biosynthesis and to become specialized for sucrose transport activity (**Figure 3A, Dataset S2**).

Taken together, the enriched GO terms of the differentially expressed genes describe different stages of morphogenesis, pointing to two trajectories of development along the leaf: development along the proximal-distal axis and the medio-lateral axis (**Figure 3B and C**). Cells that have achieved specialized photosynthetic functions, leaf development, and sugar transport define the final morphogenetic stages. Margin regions undergoing active cell division are defined by chromatin assembly and DNA processing (replication, integration, and recombination), which are required for proper cell cycle progression, while the most highly meristematic tissue in the margin region at the base is defined by only transcriptional activity and transcription factor and DNA binding (**Figure 3A-C**). Thus, the P4 tomato leaf represents a complex mixture of developmentally distinct regions that cannot be defined solely along the proximal-distal or medio-lateral axes.

### Modeling gene expression differences across the medio-lateral axis predicts that photosynthetic activity first occurs in the rachis

Differential gene expression analysis in each region along the proximal-distal axis revealed specific genes and enriched GO terms that are unique to the top, middle, or base of the leaf. Next, we tried to identify gene activity that defines rachis and margin identity across the entire P4 leaf primordium regardless of position on the longitudinal axis. To address this issue, we performed differential gene expression analysis across the margin and rachis tissue and adjusted for variability between the proximal-distal axis by employing an additive linear model using the top, middle, and base identities as a blocking factor in our experimental design using EdgeR (Robinson et al., 2010). In the wild type, across the entire proximal-distal axis, 1,089 genes were significantly up-regulated in the rachis and 188 genes were significantly up-regulated in the margin (**Figure 4A, Dataset S3**). GO enrichment analysis of these upregulated genes revealed 24 enriched GO terms in the rachis (**Dataset S4**). These terms were categorized into eight major categories: Sugar biosynthesis and transport, Metabolism (carbohydrate and glucose), Transmembrane transport, Leaf development, Photosynthesis/light harvesting, Catalytic activity, Cell wall organization, and Response to light (**Figure 4B**). These categories reflect the activities of genes that were up-regulated in the rachis compared to the margin across the entire proximal-distal axis. These results suggest that the rachis region of a P4 leaf has many specialized tissue types and may already be physiologically active.

**Figure 4.**
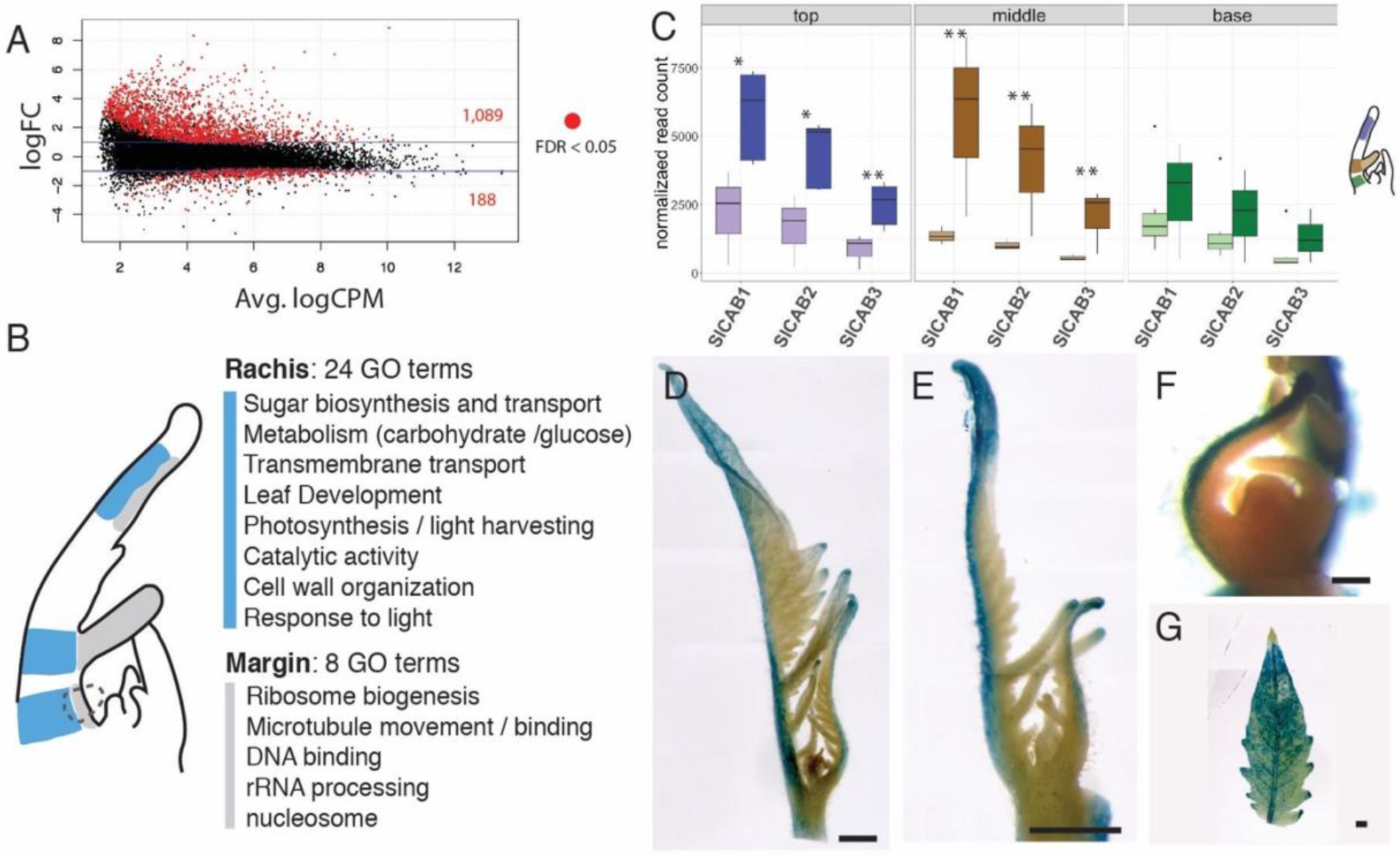
Differential gene expression between the margin and rachis in a wild-type P4 leaf and chlorophyll A-B binding gene activity in the rachis compared to margin tissue during early leaf development. (A) Results of differential gene expression analysis in the wild type showing average log Counts Per Million (LogCPM) over log Fold Change (logFC) values. The number of significant differentially regulated genes (red) between margin and rachis tissue is shown. (B) Summary of GO terms describing up-regulated genes in each tissue, showing that rachis (blue) tissue is predominantly described by GO terms related to cell specialization compared to margin tissue (gray). (C) Normalized read count for chlorophyll A-B (CAB) binding genes in tomato (*SlCAB*). Colors highlight the separation of the margin (lighter colors) and rachis (darker colors) along the top (purple), middle (brown), and base (green). (D-G) pCAB::GUS expression showing photosynthetic activity during leaf development in tomato. pCAB::GUS is localized to the rachis of P4-P6 leaflets, illustrating differential regulation of *CAB* expression along the medio-lateral axis during early leaf development. pCAB::GUS is nearly ubiquitous in (G) P8 terminal leaflet. * p-value < 0.005 for significantly up-regulated genes in rachis tissue compared to the margin based on modeled differential expression analysis. Scale bars (D-E, G) = 1 mm, (F) = 100 μm.

### Verifying photosynthetic gene expression patterns

Of the gene expression patterns described above, the most prominent pattern revealed by both pairwise and modeled differential expression analyses was the persistent presence of genes associated with GO terms related to photosynthetic processes; these genes were up-regulated in the rachis compared to margin tissues (**Figure 3 and 4**). While the up-regulation of genes involved in cell wall development, leaf development, and transport might be expected in the rachis, a region of the leaf that acts as a connective corridor to the rest of the plant, we were surprised to find up-regulation of so many genes defined by GO terms related to photosynthesis. As noted in the pairwise differential gene expression analysis described above, the most abundant enriched GO terms for up-regulated genes in the rachis were related to sugar biosynthesis and photosynthesis, indicating that the rachis region likely has functioning photosynthetic machinery before it is acquired by the P4 margin, which is destined to become the blade, the primary photosynthetic tissue of the leaf. Since little is known about when photosynthesis first begins in a developing leaf, and no previous studies have described photosynthesis specifically in the rachis, we attempted to verify the notion that the rachis is a photosynthetic force during early leaf development.

To verify the photosynthetic signature repeatedly found to be up-regulated in rachis compared to margin tissue, we searched for photosynthetic genes in our dataset that showed significant differential expression between the rachis and margin in each longitudinal region. We identified three *Light Harvesting Chlorophyll A-B binding* (*CAB*) genes (Solyc03g005760 [*SlCAB1*], Solyc03g005760 [*SlCAB2*], and Solyc03g005760 [*SlCAB3*]) with significantly up-regulated expression in the rachis regions compared to the margin (**Figure 4C, Data S1**). CAB proteins balance excitation energy between Photosystem I and II during photosynthesis (Liu and Shen, 2004) and are important components of photosynthesis.

In an attempt to verify the gene expression differences identified in our experimental set-up and visualize when and where photosynthetic activity begins in a leaf primordium, we constructed a transgenic line expressing a representative *CAB* gene promoter attached to the β-glucuronidase (GUS) reporter (*pCAB1::CAB1::GUS*) (Mitra et al., 2009; Tindamanyire et al., 2013). In the expanded leaflets of P8 leaves, *pCAB::CAB1::GUS* expression was nearly ubiquitous across the entire blade (**Figure 4D**). At this age, the leaf had the anatomy of a fully functional photosynthetic organ. As predicted from our gene expression analysis, in younger leaf primordia, we found a clear *pCAB::CAB1::GUS* signal localized predominantly in the rachis region along the proximal-distal axis in P4-P7 leaves (**Figure 4E and F**). The *pCAB::CAB1::GUS* signal spread to the distal tips of newly established leaflets and lobes after P4 and continued to spread to the margin tissue as development proceeded until the entire leaf showed expression (**Figure 4D-G**).

Since pCAB::CAB1::GUS is predominantly expressed in the rachis region during early development, we suggest that the rachis is the first region in a developing leaf to function photosynthetically, as predicted in our RNA-seq analysis. Previous studies have suggested that chloroplast retrograde signaling and sugar trigger leaf differentiation (Andriankaja et al., 2012; Lastdrager et al., 2014). In the light of these studies, the enrichment of photosynthetic genes in the rachis provides the first line of evidence that during a very early developmental stage (P4), the rachis region does not simply function as a conduit for nutrients and water transport, but it also functions in photosynthesis and sugar production.

### Self Organizing Maps (SOM) identify specific groups of genes that share similar expression patterns

To refine our results and identify groups of genes sharing similar co-expression patterns that may be too complex to define by differential expression analysis alone, we used Self Organizing Mapping to cluster genes based on gene expression patterns across the six tissue groups. SOM (Tamayo et al., 1999) begins by randomly assigning a gene to a cluster. Genes are subsequently assigned to clusters based on similar expression patterns via a reiterative process informed by previous cluster assignments. This clustering method allows genes to be grouped based on specific expression patterns shared across different tissues, allowing genes to be classified into smaller groups than those generated by differential expression analysis alone. SOM analysis also allowed us to survey the most prominent types of gene expression patterns found in our data.

To focus on the most variable genes across tissues, we selected the top 25% of genes with the highest coefficients of variation, resulting in a dataset of 6,582 unique genes (**Dataset S5**). We used principal component analysis to visualize groups of genes and found that the first four principal components explained 31.9%, 26.2%, 19.0%, and 13.5% of the amount of variation in the dataset, respectively (**S5 Figure A**). Looking at the expression of these genes in the PC space revealed distinct clusters of genes with related expression patterns (**Figure 5A**).

**Figure 5.**
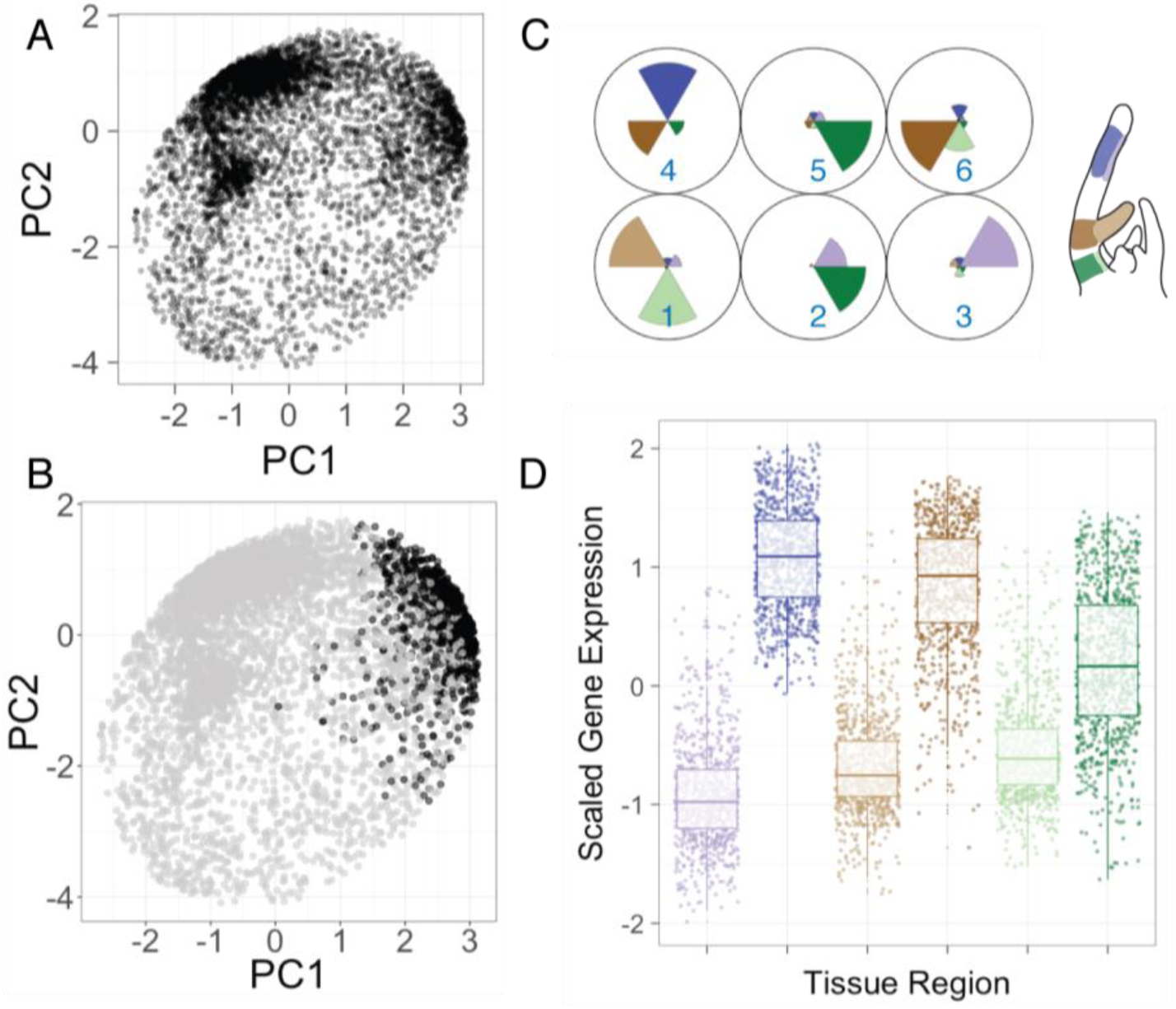
Top clusters identified by SOM analysis define each tissue region based on up-regulated genes. (A) Plotting of wild-type gene expression observed in the top 25% of genes based on coefficient of variation in the Principal Component (PC) space. (B) Projection of SOM cluster 4 onto the PC space explains one of the main clusters in the PC space (C) Codebook vector of a 2×3 SOM analysis showing the top six clusters (D) Gene expression patterns of Cluster 4 across the six tissue types. (C) and (D) Colors highlight the margin (lighter colors) and rachis (darker colors) along the top (purple), middle (brown), and base (green).

To identify the most common gene expression patterns that describe the data, SOM analysis was initially limited to six clusters (**Dataset S6**). One of these clusters, Cluster 4 (with 1090 genes), defines a clear separation of margin and rachis tissues, which again reinforces our finding that the expression levels of many genes differ depending on where the sampled tissue is localized along the medio-lateral axis (margin vs. rachis). This cluster is enriched in genes defined by Carbohydrate metabolic processes, hydrolase activity, protein dimerization, membrane, transporter activity, and photosynthesis and light harvesting (**S5 Figure, Dataset** S7). These findings mirror the results obtained by differential gene expression analysis and reflect the overall abundance and diversity of genes up-regulated in the rachis, comprising the largest signal in our dataset. These findings likely reflect the specialization that occurs in tissues as the rachis develops an identity distinct from the margin.

### Auxin transport and regulation as a defining feature of margin identity

To refine our analysis of gene expression patterns to genes that direct margin identity, we generated a larger clustering map. We used this approach to obtain a smaller subset of genes than could be obtained by differential gene expression analysis or SOM clustering using a smaller number of clusters. We were especially interested in identifying specific types of gene expression patterns that defined the medio-lateral axis; in this case, we looked for groups of genes that were preferentially up or down-regulated in the margin compared to the rachis. We specified 36 clusters in a 6×6 hexagonal topology, forcing interactions between multiple tissue types (**Figure 6A, S6 Figure**). We surveyed the gene expression patterns of each of the 36 clusters (**Dataset S8**) and identified Clusters 10 (n =108) and 11 (n=112), which describe a group of genes that were up-regulated in the margin and down-regulated in the rachis in stage P4 wild-type plants (**Figure 6B - C**). While over half of these genes (57.2%; 126/220) have no known function, many of the remaining genes are known to be involved in leaf margin identity (**Table 1**). Interestingly, Clusters 10 and 11 also contained genes related to auxin transport, biosynthesis, and regulation (*YUC4, PIN1, AUX2-11*) and genes (*ARGONAUTE7*/Solyc01g010970) known to interact with Auxin Response Factors (ARFs) (Yifhar et al. 2012).

**Figure 6.**
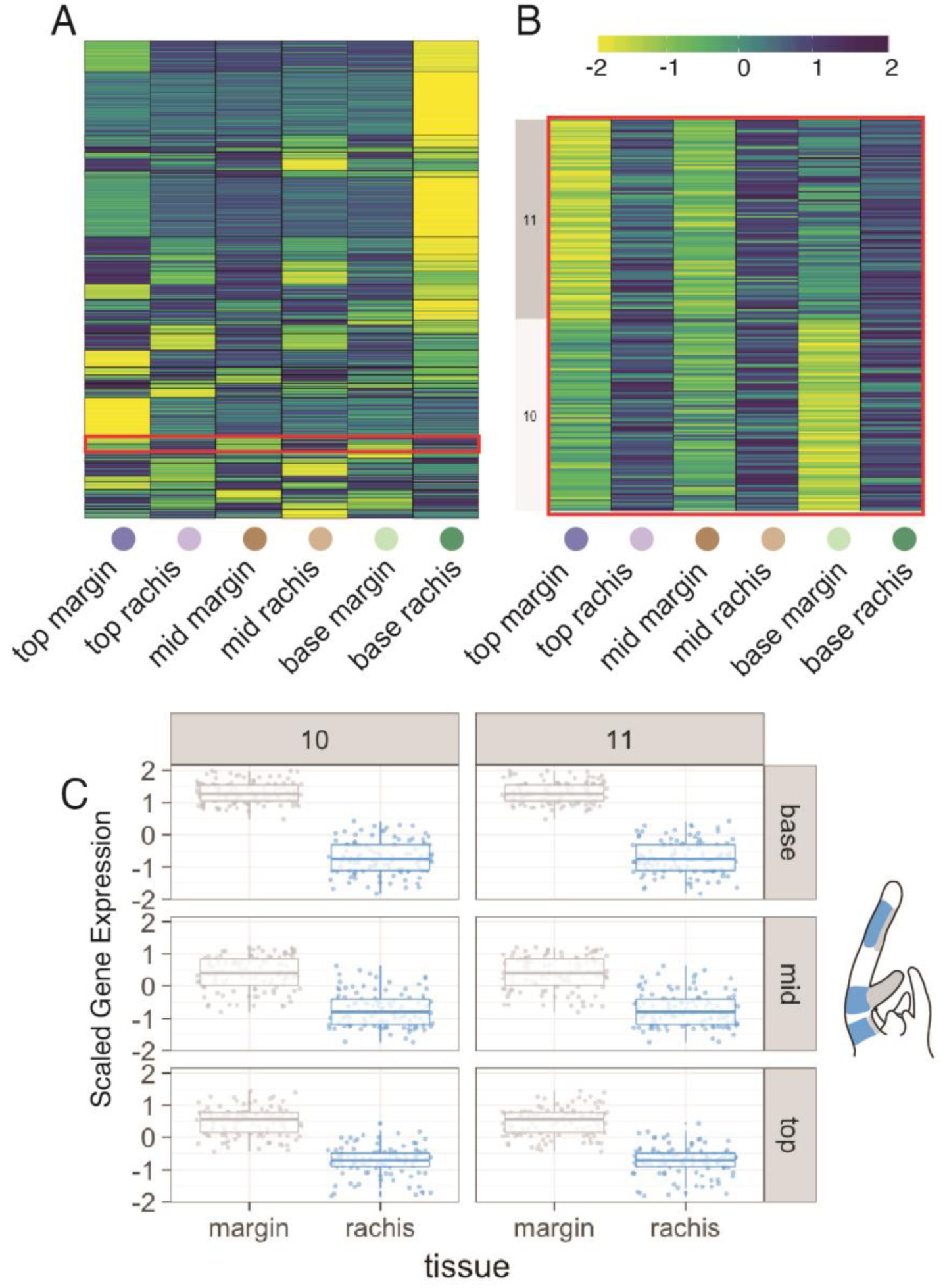
Large SOM map describes a small gene cluster that defines margin identity. (A) Heatmap representing the gene expression patterns of the 36 gene clusters. The red box highlights clusters 10 and 11 (B) Heatmap of clusters 10 and 11, with genes that are up-regulated in the margin and down-regulated in the rachis. (C) Boxplot showing the gene expression patterns of cluster 10 and 11. See S5 Figure for full heatmap of all 36 clusters.

Guided by the gene expression data in the wild type, and the results that auxin might play a role leaflet initiation in the base margin region, we wanted to see if auxin transport differences in *tf-2* could explain the striking feature of loss of meristematic potential in the basal margin of this mutant. To look specifically at the differences in auxin transport between *tf-2* and wild type and to verify the differences in *SlPIN1* gene expression found between the wild type and *tf-2*, we crossed a fluorescently labeled *pPIN1::PIN1::GFP* line (*PIN1::GFP*) (Benková et al., 2003; Koenig et al., 2009) with *tf-2* to visualize differences in PIN1 localization and expression in P4 leaves. In the wild type, PIN1::GFP was present along the entire margin region of a P4 leaf, with the strongest signal present at the site of the newly established LL1 (**Figure 7A and B**). In *tf-2*, there was an overall decrease in fluorescent signal along the margin of a P4 leaf. Also, *tf-2* had a noticeable decrease in PIN1::GFP fluorescent signal in the base margin region (**Figure 7C and D**). In addition, we visualized auxin presence using the auxin-inducible promoter DR5::Venus (Bayer et al., 2009). As observed previously (Shani et al., 2010; Martinez et al., 2016), in the wild type, DR5::Venus was expressed at the site of leaflet initiation as a sharp wedge-shaped focus region (**Figure 7E-F**). By contrast, in *tf-2*, there was a DR5::Venus focus region, but it was diffuse and located in the upper layers of the margin (**Figure 7G-H**). These results support the hypothesis that while *tf-2* is capable of forming auxin foci, it is incapable of maintaining proper auxin foci and canalization processes, as evidenced by the reduced *PIN1* expression in the basal margin region of the *tf-2* P4 leaf. The transcriptomic results and auxin visualization experiments suggest that misregulated auxin transport and biosynthesis, and specifically *SlPIN1* misregulation, are important contributors to the *tf-2* phenotype and that these processes are vital regulators of margin organogenesis.

**Figure 7.**
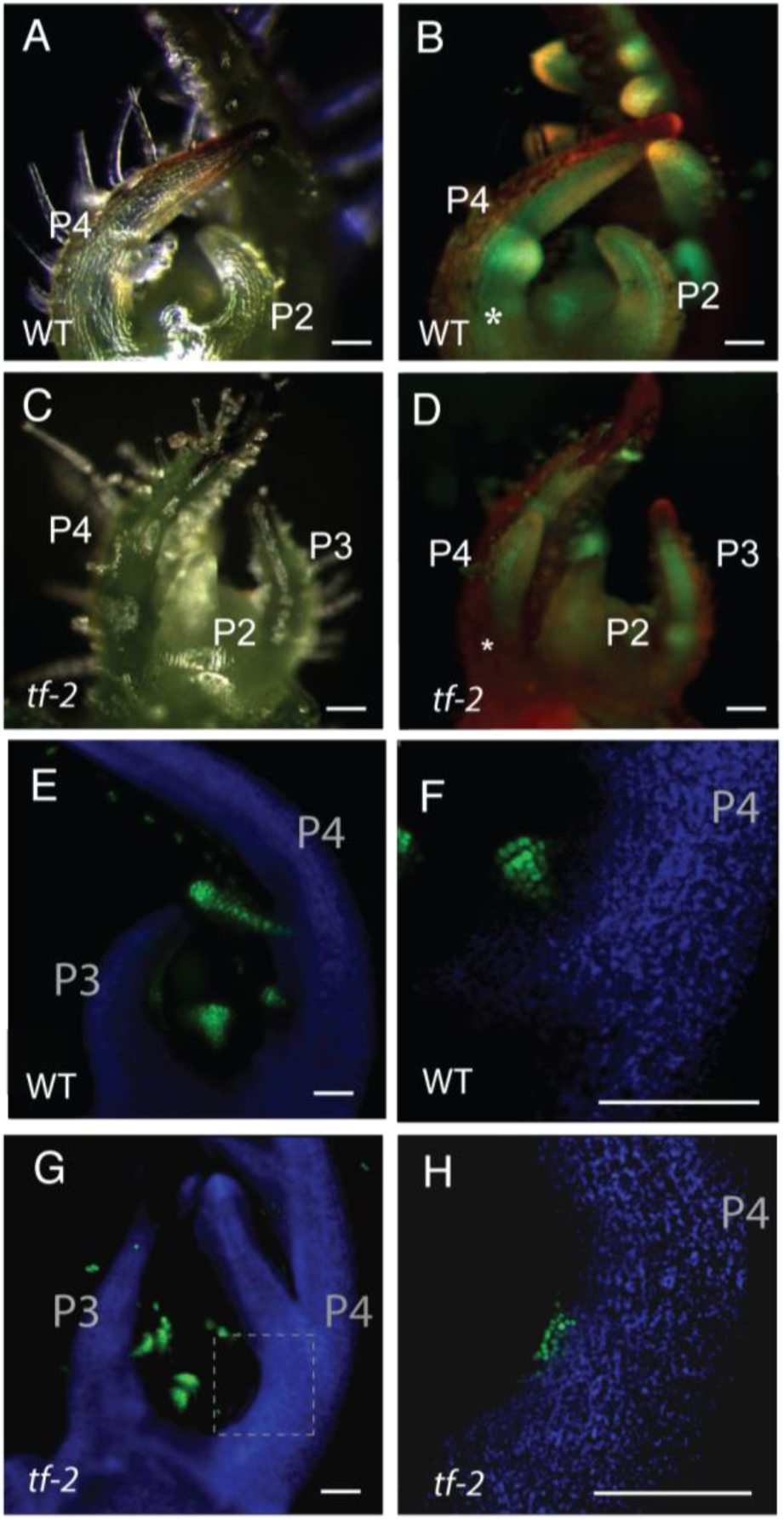
Auxin visualization during leaflet initiation in wild type and *tf-2*. (A)-(D) Microscope images of apices from (A) and (B) wild type and (C) and (D) *tf-2*. (B) and (D) Fluorescence signals of PIN1::GFP (green) and chlorophyll autofluorescence (red) * marks the base marginal blastozone region. (B) shows clear PIN1::GFP signal in wild type along the entire margin of the P4 leaf, while in (D), *tf-2* has lost signal in the base marginal blastozone region. (E)-(H) DR5::Venus signal (green) observed by confocal microscopy. (E) and (F) wild type plant apices. (G) and (H) *tf-2* plants apices (F) and (H) close up on the site of leaflet initiation of the base margin region of P4 leaves. Scale bars = 100 μm.

### Differences in gene expression patterns between the wild type and *tf-2* help define the loss of basal meristematic potential in *tf-2* leaves

We included *tf-2* in this study because it has the intriguing phenotype of being unable to form new leaflets after the first two LL1 leaflets. At the P4 stage, *tf-2* has already lost the organogenetic ability to initiate new leaflets. As revealed by our auxin transport visualization analysis, *tf-2* appears to receive a leaflet initiation signal, as it is capable of forming auxin foci (**Figure 7H**), but the tissue is unable to initiate leaflet organs. We examined our data to determine whether gene expression differences could explain the loss of meristematic competency in *tf-2*. We performed differential gene expression analysis of only *tf-2* reads. There were a lot fewer differentially expressed genes between the margin and rachis in the top, middle, and base regions of this mutant compared to the wild type (**S3 Figure A, Dataset S1**). However, *tf-2* followed similar gene expression trends to the wild type when margin and rachis identity were compared. The margin was more enriched in genes related to cell division and cell expansion, while the rachis was enriched in genes related to specialization, including water transport, metabolic processes, photosynthesis, and leaf development; however, these differences were mostly apparent in the base region of the *tf-2* mutant (**S3 Figure B**). The main difference between wild type and the *tf-2* mutant was a reduction in up-regulated differentially expressed genes in the rachis region compared to the margin in top, middle, and base regions (**S3 Figure A**). It should be noted that while wild type and *tf-2* were similar morphologically at the P4 stage, the *tf-2* mutant appeared to be further along in the morphogenesis process in all regions (top, middle, and base), a feature described by Naz and coworkers (2013). This overall difference in the two genotypes should be taken into account at the morphological level, and, as evidenced by our transcriptional analysis, at the molecular level. In the margin of *tf-2*, we examined the differentially expressed genes between the rachis and margin and found many genes related to leaf development.

Taking into account the overall differences between these two genotypes, we were interested in understanding why *tf-2* is unable to initiate lateral leaflets beyond LL1. Could transcriptional differences explain the loss of morphogenic capacity in *tf-2*? To address this issue, we combined both genotypes and used a generalized linear model (glmQLFTest in edgeR) in which we defined each genotype as a group and therefore could compare the top, middle, and base regions of the two genotypes. When we compared the base margin region between *tf-2* and the wild type (**Figure 1A**), only 23 genes were differentially expressed: all were downregulated in the wild type compared to *tf-2* (**Table 3**). We focused on the 12 genes that were functionally annotated and noticed that *SlBOP2* was significantly up-regulated in the margin of *tf-2* compared to the wild type (**Figure 8B**).

**Figure 8.**
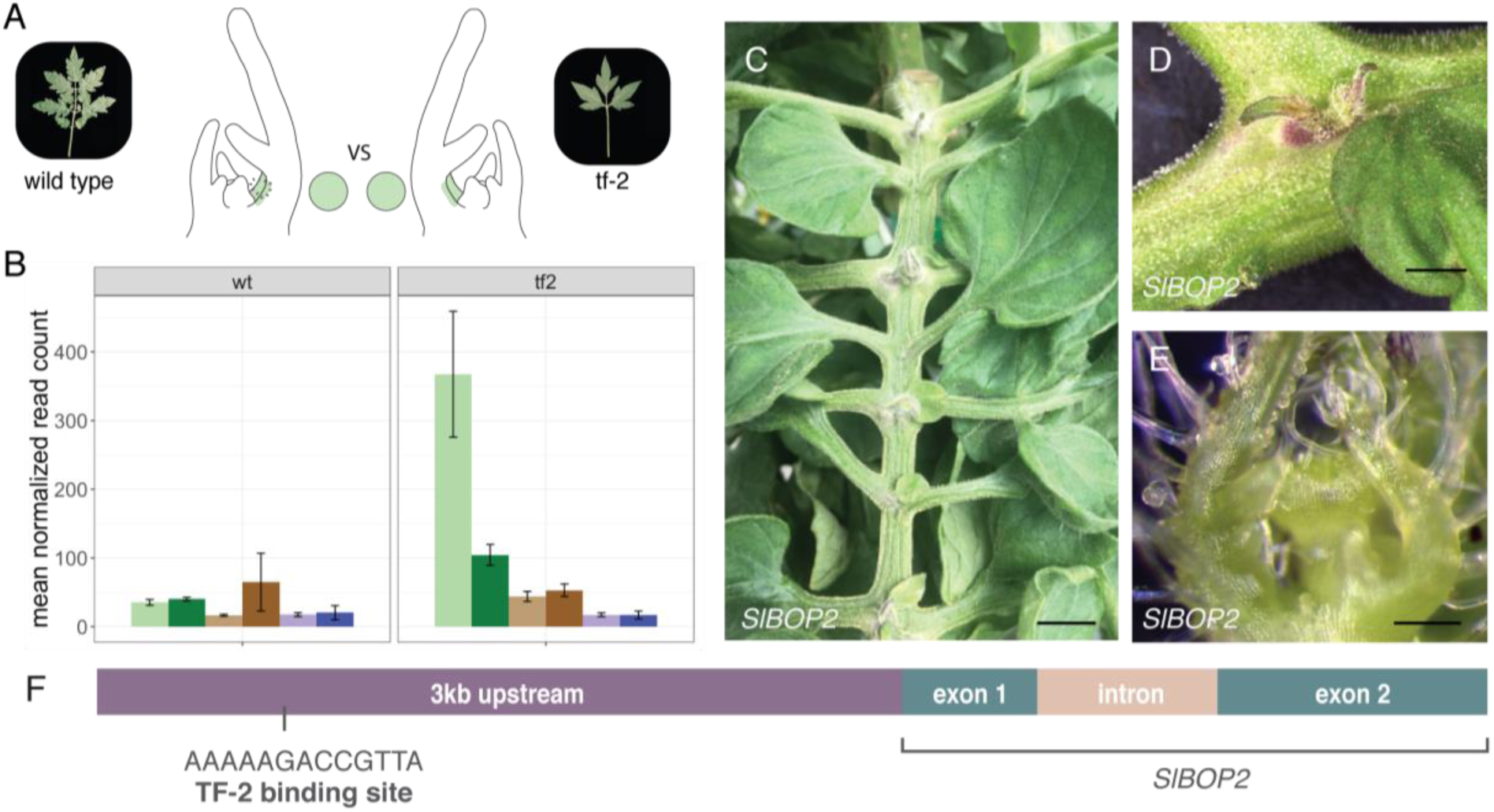
Differential gene expression analysis in base margin tissue between wild type and *tf-2* reveals BOP2 as a regulator that suppresses meristematic identity. (A) Schematic illustrating the regions (base margin) compared between wild type and *tf-2* using modeled differential gene expression analysis (B) Bar graph illustrating the expression patterns of *SlBOP2* across all six tissue types between wild type and *tf-2*, showing how *SlBOP2* is upregulated in the base margin only in *tf-2*. Colors highlight the separation of the margin (lighter colors) and rachis (darker colors) along the top (purple), middle (brown), and base (green). (C) - (E) *SlBOP2* CRISPR knockout line (*CR-slbop2*), which displays ectopic shoot apical meristems along the rachis of complex leaves. (F) *SlBOP2* genomic region. Black line shows the *TF-2* binding site 3kb upstream of *SlBOP2*. Scale bars (C) = 10 mm, (D) = 2 mm, (E) = 0.2 mm

We explored the role of *SlBOP2* in regulating margin and rachis tissue identity by phenotyping CRISPR/Cas9 gene edited loss-of-function *SlBOP2* mutants (*CR-slbop2)* (Xu et al., 2016) *with a focus on leaf phenotypes. Surprisingly, in CR-slbop2* plants, we observed ectopic meristems on mature leaves when plants were approximately two months old. The ectopic meristems occurred along the adaxial rachis of mature leaves at the bases of primary leaflets (**Figure 8C, D, E**). These ectopic SAM structures did not persist as the leave aged, appearing to undergo tissue death approximately three weeks after appearing on the rachis (S6 A) and only rarely did they generate complex leaf-like organs (**S6 C Figure**). Loss of function of *SlBOP2* also resulted in increased leaf complexity (**S6 B Figure**), as reported previously in these mutants (Xu et al., 2016) and in *SlBOP2* knockdown lines (Ichihashi et al., 2014). Since TF is a known transcription factor, we checked for TF binding site motifs in the 3KB upstream region of *BOP2* and found one TF binding site (**Figure 8F**). Taken together, these findings indicate that *SlBOP2* functions in determining margin meristematic identity along the rachis of the leaf and the possibility that *SlBOP2* functions via the direct binding of TF to its upstream regulatory region – although more validation is needed to verify this interaction.

## DISCUSSION

### Unique genetic signatures define leaf development along the proximal-distal and medio-lateral axes

The overall goal of this work was to use gene expression signatures to gain a better understanding of the processes that regulate morphogenesis along the medio-lateral axis in an early developing compound leaf. Based on anatomical analysis, we chose six unique regions in the P4 leaf (**Figure 1C-F**). We analyzed differential gene expression between margin and rachis tissue in the top, middle, and base regions and identified signature patterns of gene regulation along the proximal-distal (tip-base) axis (**Figure 3**) that help define leaf morphogenesis in the early tomato leaf primordium.

In addition to a basipetal wave of differentiation along the proximal-distal axis, the leaf differentiates from the midrib/rachis out into the margins at each region on the proximo-distal axis. These two regions, the margin and rachis, have distinct developmental trajectories: the rachis matures early and the marginal blastozone retains some meristematic potential and gives rise to the leaf blade region, as well as leaflets in compound leaves. Separating the rachis from the marginal blastozone region at three different points along the proximal-distal axis allowed us to determine whether development proceeds uniformly along the proximal-distal axis or if the leaf has a mosaic of developmental states in each segment along the proximal-distal axis. The further along in morphogenesis a region was, the more diverse the GO categories of genes that were upregulated in the region, likely because the last stage of leaf morphogenesis, cell specialization, had occurred. After summarizing the enriched GO terms in each of the three regions along the proximal-distal axis, patterns of developmentally distinct processes were identified in the rachis regions compared to other tissues (**Figure 3**). The margin regions, classically defined as the marginal blastozone or marginal meristem, retain the potential to divide and differentiate and also exhibit a basipetal gradient of gene expression changes of differentiation from the tip to the base of the leaf. Thus, this analysis suggests that defining leaf development or capturing gene expression in the entire primordium, or even in regions along the proximal-distal axis, does not provide an accurate picture of developmental patterns in a leaf. Further dissecting these events at cellular resolution should help define these patterns even more accurately.

### Photosynthetic capability in the rachis as a regulator of medio-lateral differentiation

To further define rachis and margin identity, we fitted an additive model that adjusts differential gene expression comparisons based on baseline differences that occur between the margin and rachis. We then performed differential gene expression analysis to reveal gene expression trends that define margin and rachis tissue regardless of the position on the proximal-distal axis. The most prevalent, though unexpected, gene expression signature we observed was the enrichment of genes associated with photosynthesis in the rachis, which we found by both differential expression analysis (**Figure 3 and 7**) and cluster analysis (**Figure 5**). Since little is known about when photosynthetic capacity is acquired during early leaf morphogenesis, we further verified photosynthesis activity using a CAB::GUS reporter (**Figure 4C-G**). This analysis suggested that photosynthetic activity is acquired as early as P4 and is not uniformly distributed along the proximal-distal and medio-lateral axes. When viewed in the context of cell differentiation processes along each axis, it is not surprising that specialized functions are first acquired in regions that mature earliest, although the function of photosynthesis has been traditionally assigned to the blade. What are the developmental consequences of sugar biosynthesis in the rachis during early leaf organogenesis? Could the rachis be the source of morphogenic signaling towards the more immature base along the proximal-distal axis and along the medio-lateral axis to the margin? Multiple studies in *A. thaliana* identified thousands of genes that respond to changes in sugar levels by modifying transcript abundance (Price et al., 2004; Bläsing et al., 2005; Osuna et al., 2007; Usadel et al., 2008). These studies suggest that the main photosynthetic product, sugar, functions as a signal for plant development and growth.

In the P4 primordium under study, while the rachis has acquired specialized functions, the margin is actively dividing, a process that relies on cell cycle progression. The cyclin genes *CYCD2* and *CYCD3*, encoding critical regulators of the cell cycle, are up-regulated in response to sugar (Riou-Khamlichi et al., 2000). Interestingly, sucrose has also been shown to influence auxin levels (Lilley et al., 2012; Sairanen et al., 2012), transport, and signal transduction (Stokes et al., 2013), and metabolism (Ljung, 2013). Moreover, sugar accumulation is spatiotemporally regulated in meristematic tissue in both the shoot and root apical meristem (Francis and Halford, 2006). Is the development of photosynthetic capacity in the rachis a cause or a consequence of its early differentiation? Do the acquisition of photosynthetic capability and the production of sugars represent a global mechanism for signaling quiescent regions to progress into the cell division phase? More work exploring photosynthesis, sugar transport, hormone regulation, and gene expression should help uncover a possible role for the rachis in regulating morphogenetic processes during early leaf organogenesis.

### The presence of auxin as a defining feature of organogenic potential in margin tissue

PIN1-directed auxin transport is an important regulator of leaf development (Reinhardt et al., 2003; Heisler et al., 2005; Scarpella and Helariutta, 2010; Kawamura et al., 2010; Hay and Tsiantis, 2006; Scarpella et al., 2006) and leaflet initiation (Koenig et al., 2009). A common mechanism unites PIN1-directed development during leaf organogenesis across the systems studied: PIN1 directs auxin along the epidermal layer to sites of convergence on the meristem and transports the auxin subepidermally into the internal layers (Scarpella et al., 2010). *PIN1* can be split into two highly supported sister clades: *PIN1* and *Sister of PIN1* (*SoPIN1*) (O’Connor et al., 2014; Bennett et al., 2014; Abraham Juárez et al., 2015). The *SoPIN1* and *PIN1* clades might have disparate but complementary functions in auxin transport during organ initiation, where SoPIN1 mainly functions in epidermal auxin flux to establish organ initiation sites and PIN1 functions in the transport of auxin inward (O’Connor et al., 2014; Abraham Juárez et al., 2015; Martinez et al., 2016). Tomato has one gene in the *PIN1* clade (*SlPIN1*) and two genes in the *SoPIN1* clade (*SlSoPIN1a* [Solyc10g078370] and *SlSoPIN1b* [Solyc10g080880]) (Pattison and Catalá, 2012; Nishio et al., 2010; Martinez et al., 2016). The current findings suggest that in *tf-2, SlPIN1* is down-regulated at the region of leaflet initiation compared to the wild type. Using *PIN1::GFP* as a reporter, we observed a lack of fluorescence in the base marginal blastozone region of *tf-2* (**Figure 7C-G**). Using DR5:VENUS as a reporter, we observed the diffuse localization of auxin in the base margin region of *tf-2* apices. Interestingly, even with external auxin application, *tf-2* is not capable of leaflet initiation (Naz et al., 2013), suggesting that the ability to direct auxin inwards using PIN1, and not auxin accumulation itself, may be compromised in this mutant. It remains to be seen whether a common theme for organ formation will emerge in organisms in which the *PIN1* clade has diverged into two groups. Based on our findings about the marginal blastozone region in *tf-2*, we suggest that in addition to the creation of auxin foci, the drainage of auxin into internal leaf layers might also be required for leaflet initiation. Analysis of higher-order mutants in the larger *PIN1* clade should help resolve this issue.

### Organogenic potential of the margin and homology of the leaf margin and the SAM

In the current study, we were especially interested in obtaining a genetic understanding of the loss of organogenetic potential in the base margin of *tf-2*. We specifically looked for the transcriptional differences that explain the loss of organogenetic potential in *tf-2* compared with the wild type, specifically in the margin base region. This led us to a small list of 23 differentially expressed genes including *SlBOP2* (**Figure 8, Table 2**). Characterization of the *CR-bop2* line (**Figure 8**) revealed ectopic SAM production along the adaxial rachis at the bases of primary leaflets of the complex leaf. This finding supports the notion that the suppression of meristematic identity by BOP family members is important during leaf morphogenesis. BOP1 was first introduced as a suppressor of lamina differentiation on the petioles of simple *Arabidopsis* leaves (Ha et al., 2003, 2004) that limits meristematic cell activity, as the *bop1* mutant displays ectopic meristematic cells beyond the boundary between the base of the blade and petiole (Ha et al., 2003, 2004). Further work using *SlBOP* knockdown and knockout tomato lines demonstrated that SlBOP2 suppresses organogenetic potential (S6 Figure; (Xu et al., 2016; Ichihashi et al., 2014), as lines with reduced or absent *SlBOP* function showed increased leaflet organ initiation/leaf complexity. BOPs interact with transcription factors to regulate floral identity, including the interaction of BOP with PERIANTHIA (PAN) in Arabidopsis (Hepworth and Pautot, 2015) and the interaction of TERMINATING FLOWER (TMF) with *SlBOPs* to repress meristematic maturation in tomato flowers (MacAlister et al., 2012; Xu et al., 2016). We further hypothesize that *SlBOP2* and a transcription factor interact to regulate organogenic potential in complex leaves. Specifically, perhaps the TF transcription factor binds to the upstream regulatory region of *SlBOP2* (**Figure 8F**). We suggest that both *TF and SlBOP2* function in suppressing meristematic properties of the margin during an early developmental window that gradually closes with leaf maturation, an idea consistent with the view of the marginal blastozone described by Hagemann (1970).

While our understanding of the recruitment of genetic regulators in a spatiotemporal context continues to increase, one of the more exciting questions still remains: Is the marginal meristem evolutionarily derived from the SAM (Floyd and Bowman, 2010)? Ectopic adventitious SAMs have been shown to occur on leaves of functional knockouts of *CUP-SHAPED-COTYLEDONS2* (*CUC2*) and *CUC3* (Blein et al., 2008; Aichinger et *al., 2012*; Hibara et al., 2003) and of Arabidopsis lines overexpressing homeobox genes *KNOTTED-1 (KN1)* and *Kn1-like* (*KNAT1*) (Chuck et al., 1996; Sinha and Hake, 1994) in a region analogous to the base of an emerging leaflet, suggesting developmental analogy and possibly homology to axillary meristems. Axillary meristems form on the adaxial surface at the boundary zone between the leaf and SAM, where BOP2 has already been shown to play a regulatory role in this process in tomato (Izhaki et al., 2018), barley (*Hordeum vulgare*) (Tavakol et al., 2015; Dong et al., 2017), and maize (Dong et al., 2017). The ectopic meristem phenotype of *CR-slbop2* on the margins of complex tomato leaves suggests that signals might be recruited in the margin that are similar to those present in leaf initiation sites during axillary meristem formation. Our findings add further evidence that the margin is analogous, and possibly homologous at the process level, to the SAM. The leaf margin likely evolved via the genetic recruitment of similar regulatory factors, including BOP, reinforcing the importance of the reiteration of genetic mechanisms to establish distinct spatial identities in neighboring domains during plant development.

Our current understanding of the leaf margin is based on foundational work that defined the margin by explicitly tracking developmental landmarks (Avery, 1933; Poethig and Sussex, 1985a; Dolan and Poethig, 1998; Wolf et al., 1986). Early literature defined the leaf primordium as broadly meristematic during early development, with this meristematic potential becoming restricted and gradually lost as the leaf develops (Foster, 1936; Hagemann and Gleissberg, 1996; Sachs, 1969). Although such studies provide a roadmap for describing growth patterns in the margin, a major challenge is to understand how these patterns are specified at the genetic level (Whitewoods and Coen, 2017; Coen et al., 2017) and how this fits with our interpretation of the recruitment of regulatory mechanisms suppressing the morphogenetic potential of the margin during the evolution of leaves in seed plants. Plant development is reliant on reiterative patterning, and leaf development is no exception. Our findings suggest that we can describe leaf development as the reiteration and modulation of similar evolutionarily derived genetic programs that act to suppress the morphogenetic and organogenetic potential of meristematic regions in order to achieve final leaf form. Follow-up studies in additional species are needed to understand these evolutionarily conserved mechanisms and how they have been modulated to sculpt the diversity of leaf forms observed in nature.

## METHODS

### Plant growth and tissue embedding

Seeds of tomato mutant *tf-2* (LA0512) and wild-type line Condine Red (LA0533) were obtained from the Tomato Genetics Resource Center (TGRC). The seeds were sterilized with 50% bleach for two minutes and rinsed 10 times with distilled water. The seeds were placed on moist paper towels in Phytotrays (Sigma-Aldrich), incubated in the dark for two days, and allowed to germinate in a growth chamber for four days before being transferred to soil for 8 days of growth; the seedlings were grown for a total of 14 days. Generation of the transgenic DR5::Venus (cv. M82) line was described in (Shani et al., 2010) and the *AtpPIN1::PIN1::GFP* (cv. Moneymaker) line was described in (Bayer et al., 2009).

CR-bop-2 RNAi lines were received from the Lippman Lab at Cold Spring Harbor. CR-bop-2 RNAi and wild type plant (M82) were germinated and grown in growth chambers for two weeks following methods above. Plants were further grown in a growth chamber for one month. After one month, plants were transferred to a greenhouse and grown under normal light conditions. The plants were observed, and any abnormalities were noted, dated, and photographed throughout the lifetime of each plant. The images are taken from mature leaves from plants that were 63 days old. Leaf complexity counts (S6 Figure) were taken at age 45 days old.

Plants were collected in the afternoon, vacuum infiltrated for one hour with ice-cold 3:1 (100% EtOH: 100% acetic acid) fixative, and fixed overnight at 4**°**C. The samples were washed three times in 75% EtOH, passed through an EtOH series on a shaker at room temperature for one hour per step (75%, 85%, 95%, 100%, 100%, 100%), and incubated in 100% EtOH overnight at 4°C. All ethanol solutions were made using 2X autoclaved diethylpyrocarbonate (DEPC) treated water. The tissue was passed through a xylene/EtOH series for two hours per step (25%, 50%, 75%, 100%, 100%) on a shaker at room temperature. The tissue was incubated overnight at room temperature in 100% xylene with 20-40 paraffin chips (Paraplast x-tra, Thermo Fisher Scientific), followed by incubation at 42**°**C until the paraffin dissolved. The paraffin:xylene solution was subsequently removed and replaced with 100% paraffin and the sample was incubated for 3 days at 55**°**C, with the solution changed twice daily. The tissue was then embedded using tools and surfaces that had been washed with RNAseZap (Thermo Fisher Scientific) and DEPC. The embedded blocks were transversely sectioned at 5 to μm thickness using a Leica RM2125RT rotary microtome (Leica Microsystems) on RNase-free polyethylene naphthalate PEN membrane slides (Leica). The slides were dried at room temperature and deparaffinized with 100% xylene.

### EdU Visualization

Cell division was visualized by observing fluorescent signals derived from an EdU incorporation assay in which EdU is incorporated into cells during the S-phase (Kotogány et al., 2010). The EdU assay was performed as previously described (Nakayama et al., 2014; Ichihashi et al., 2011) with some modifications using a Click-iT ® EdU Alexa Fluor® Imaging kit (Invitrogen). Fourteen-day-old seedlings were dissected under a microscope. After removing older leaves, P4 leaf epidermis was nicked using an insect mounting needle to increase infiltration in subsequent steps. The plant apex was incubated in water containing 10 μM EdU for two hours. The samples were washed in 1x phosphate-buffered saline solution (PBS, pH 7.4) and fixed in FAA under vacuum infiltration for 3 h. Subsequently, the samples were fixed in 3.7% formaldehyde in PBS (pH 7.4) for 30 min and washed three times in PBS with shaking. Alexa Fluor coupling to EdU was performed in the dark following the manufacturer’s instructions. Photographs were taken under a Zeiss LSM 710 Confocal Microscope with excitation wavelengths set at 488 and 420 nm.

### Flow Cytometry and GUS Staining

Ploidy levels were measured using a PA-I ploidy analyzer (Partec) as described previously (Sugimoto-Shirasu et al., 2002). We grew up 50 plants and sampled the oldest two leaves on the plant at each timepoint. To identify the leaves by counting from youngest leaves, requires apical meristem destruction, therefore we choose to use the count Leaf age from oldest leaf -Leaf 1 corresponds to the oldest leaf, Leaf 2 is second oldest and so on. For the first two timepoints (Day 8 and 18) we sampled L1 or L2 and for the later timepoints, Day 30 and above, we sampled the oldest intact leaf, which because of age was often damaged, and therefore the leaves sampled ranged from Leaf 2 – Leaf 5. All leaves sampled beyond Day 30 had reached maturity. Fresh tissue was extracted from whole leaves at the youngest leaf age (Day 8), while older stage tissue was extracted from both the top and bottom sections of the leaf (Day 18 – 90). The tissue was chopped with a razor blade. Cystain extraction buffer (Partec) was used to release nuclei. The solution was filtered through a CellTrics filter (Partec) and stained with Cystain fluorescent buffer (Partec). At least 4000 isolated nuclei isolated were used for each ploidy measurement. Flow cytometry experiments were repeated at least three times using independent biological replicates.

Histochemical localization of GUS activity was performed as previously described (Kang and Dengler, 2002). Representative images were chosen from >15 samples stained in three independent experiments.

### Laser Capture Microdissection and RNA processing

Each tissue type was independently captured through serial sections using a Leica LMD6000 Laser Microdissection System (Leica Microsystems). Each biological replicate contained tissue collected from 5-8 apices. S1 Figure and Movie 1 show how tissue regions were identified and dissected. Tissue was collected in lysis buffer from an RNAqueous®-Micro Total RNA Isolation Kit (Ambion) and immediately stored at -80 °C. RNA extraction was performed using an RNAqueous®-Micro Total RNA Isolation Kit (Ambion) following the manufacturer’s instructions. The RNA was amplified using the WT-Ovation™ Pico RNA Amplification System (ver. 1.0, NuGEN Technologies Inc.). The RNA was purified using RNAClean ® magnetic beads (Agencourt) and processed within one month of fixation to ensure RNA quality.

RNA-seq libraries were created as described by Kumar and coworkers (Kumar et al., 2012), starting with the second-strand synthesis step, with the following modifications: For second-strand synthesis, 10 µL of cDNA (>250 ng) was combined with 0.5 µl of random primers and 0.5 µl of dNTP. The sample was incubated at 80°C for 2 min, followed by 60°C for 10 sec, 50°C for 10 sec, 40°C for 10 sec, 30°C for 10 sec, and 4°C for at least 2-5 min. After adding 5 µl of 10x DNA pol buffer, 31 µL water, and 2.5 µL DNA Pol I on ice, the sample was incubated at 16°C for 2.5 h. The process was continued following the published (Kumar et al., 2012) protocol starting with step 2.3: Bead purification of double-stranded DNA. The libraries were quality checked and quantified using a Bioanalyzer 2100 (Agilent) on RNA 6000 Pico Kit (Agilent) chips at the UC Davis Genome Center. The libraries were sequenced in three lanes using the HiSeq2000 Illumina Sequencer at the Vincent J Coates Genomics Sequencing Laboratory at UC Berkeley.

### Read Processing, Differential Expression, and Gene Ontology (GO) Enrichment analysis

Quality filtering, N removal, and adaptor trimming were performed on data from each of the three Illumina sequencing lanes separately. We first performed N removal using read_N_remover.py. Sequences below a quality (phred) score of 20 were removed without reducing the read size to below 35 bp. To remove adapter contamination, we used adapterEffectRemover.py, setting the minimum read length to 41. To assess the quality of the reads after pre-processing, we ran FASTQC (available at http://www.bioinformatics.bbsrc.ac.uk/projects/fastqc/) before and after pre-processing. To filter out reads from chloroplast or mitochondrial sequences, all libraries were mapped to the S.lycopersicum_AFYB01.1 mitochondrial sequence from NCBI and the NC_007898.3 chloroplast sequence from NCBI using STAR 2.4.0 (Dobin et al., 2013). Reads that did not map to either organelle were mapped to the ITAG3.10 *Solanum lycopersicum* genome using STAR 2.4.0, where non-genic sequences were masked using the inverse coordinates of the ITAG3.10 gene model gff file. Bedtools (Quinlan, 2014) coverageBed was then used to count mapped reads, using a bed file generated from ITAG3.10 gene models. We built an online visualization tool for the community to manually explore the reads generated across the six tissue types in both wild type and *tf-2*: bit.ly/2kkxsFQ (Winter et al., 2007; Patel et al., 2012).

Read processing and differential expression analysis were performed using the R package edgeR (Robinson et al., 2010). Pairwise differential gene expression in each region along the proximal-distal axis was calculated in each proximal-distal region (top, middle, base) in separate analyses. Differential gene expression was determined using ‘exactTest()‘, multiple testing correction was performed using the Benjamini–Hochberg procedure, and significance of differential expression was determined using a cutoff of FDR < 0.05. To estimate differential expression of genes across the entire marginal blastozone and rachis regions, we used an additive linear model where the proximal-distal axis was assigned as a blocking factor, which adjusts for differences between the margin and rachis in the top, middle, and base: model.matrix(∼Region + Tissue). For both pairwise and modeling analysis of differential expression, counts per million were calculated from raw reads, and genes with < 5 reads in 2 or more reps were removed. We estimated common negative binomial dispersion and normalized counts across all samples using the trimmed mean of M-value (TMM) method (Robinson and Oshlack, 2010). Normalized Read counts, as calculated by Counts Per Million (cpm), are available in **Dataset S9**. GO enrichment analysis was performed using the R libraries GO.seq and GO.db. The GO terms were summarized using REVIGO (http://revigo.irb.hr/). The full code used for these analyses is available at http://github/iamciera/lcmProject (DOI provided upon publication).

### SOM clustering

To explore the genes whose expression levels were the most variable across tissues, we identified the top 25% genes based on coefficient of variation and ratio of standard deviation compared to mean from our count data. To remove differences in counts between samples, the magnitude of gene expression data was scaled between 2 and -2 in wild type and *tf-2* separately using the ‘scale()’ function (Team, 2014). Hexagonal layout was used for all SOM clustering (Kohonen). For basic SOM analysis, the SOM() function was used for each genotype separately, while superSOMs were performed using superSOM() in the Kohonen R package (Wehrens and Buydens, 2007). Training for both methods was performed in 100 iterations in which the adaptive learning rate decreased from 0.05 to 0.01. Codebook vectors and distance plots of cluster assignments were generated using the visualization functions in the Kohonen R package (Wehrens and Buydens, 2007) and ggplot2 (Wickham, 2009). To ensure that the major variances in gene expression patterns were defined by SOM clustering and to verify consistency in clustering, cluster assignments were projects onto the PC space. All scripts used in clustering are available at http://github/iamciera/lcmProject (DOI provided upon publication).

## AUTHOR CONTRIBUTIONS

Conceived and designed the experiments: CM, NS. Performed molecular experiments and plant characterizations: CM. Plant maintenance and phenotyping of *CR-bop2* lines: SL. Contributed plant lines, protocols, and/or reagents: NS, KS. Performed computational analysis: CM. Performed read mapping: MW. Analyzed the data: CM, NS, KS. Wrote the paper: CM, NS. Edited the paper: CM, NS, KS.

## ACKNOWLEDGEMENTS

The UC Davis Tomato Genetics Resource Center provided tomato germplasm, AtpPIN1: PIN1: GFP and the DR5: VENUSx6 lines are gifts from Cris Kuhlemeier (University of Bern) and Naomi Ori (Hebrew University, Israel), respectively. We thank Zachary Lippman (Cold Spring Harbor Laboratory) for the *CR-slbop2* lines. We would also like to acknowledge Eddi Esteban, Asher Pasha, and Nicholas J. Provart for building the EFP browser. C.C.M. was supported by a National Science Foundation Graduate Research Fellowship (DGE-1148897), Katherine Esau Summer Fellowship, Walter R. and Roselinde H. Russell Fellowship, and Elsie Taylor Stocking Fellowship. Part of the work was supported by NSF PGRP grants IOS–0820854 (to N.R. S., Julin Maloof and Jie Peng), and S.L was partially supported by IOS 1856749 (to Julia Bailey-Serres, Siobhan Brady, Roger Deal, Uta Paszcowsky, and N. R. S.). C.C.M. was also supported by a collaboration between the National Science Foundation and the Japan Society for the Promotion of Science with a Graduate Research Opportunities Worldwide (GROW) award. This material utilized resources supported by the National Science Foundation under Award Numbers DBI-0735191, DBI-1265383, and DBI-1743442. URL: www.cyverse.org. We thank Daniel Chitwood (Michigan State University) who provided computational training and insightful discussion which greatly assisted the research and Kristina Zumstein (UC Davis) for her work and organization of several experiments. We would also like to thank Siobhan Brady (UC Davis) and Andrew Groover (USDA) for their helpful comments and John Harada (UC Davis) and Julie Pelletier (UC Davis) for use and training on the laser capture microdissection microscope.

## Supplemental Data

**S1_Dataset_allsig_DE_seperately.csv** - Results of differential gene expression analysis between the margin and rachis in the top, middle, and base regions of both genotypes (WT and *tf-2*).

**S2_Dataset_sig_go_terms.csv -** GO terms describing differentially expressed genes between the margin and rachis in the top, middle, and base regions of both genotypes (WT and *tf-2*).

**S3_Dataset_wt_modelled_DE.txt -** Results of differential expression analysis across the margin and rachis tissues performed with only wild-type reads and adjusted for variability between the proximal-distal axis. An additive linear model was employed using the top, middle, and base identities as a blocking factor in our experimental design.

**S5_Dataset_top25_coefficent_of_variation.csv -** Genes with the most variable expression. The top 25% of genes were selected based on coefficient of variation, resulting in a dataset of 6,582 unique genes.

**S6_Dataset_wt_SOM_small_cluster_assignments.csv -** SOM cluster assignments for wild type using a codemap vector of 6 showing the top six gene expression clusters.

**S7_wt_SOM_small_sigGOterms.csv -** GO terms derived from Dataset S6.

**S8_wt_SOM_large_cluster_assignments.csv** - SOM cluster analysis using a codemap vector of 36 in wild type.

**S9_Dataset_normalizedReadCount_cpm.csv** - Normalized Read counts calculated as counts per million (cpm).

**S10_Dataset_tf2_modelled_DE.txt -** Results of differential expression analysis across the margin and rachis tissue performed with only *tf-2* reads and adjusted for variability between the proximal-distal axis. An additive linear model was employed using the top, middle, and base identities as a blocking factor in our experimental design.

## Notes

https://github.com/iamciera/lcmProject

### Competing Interest Statement

The authors have declared no competing interest.

### Summary of Updates

Editing the general flow and grammar. And minor updates to interpretation of results. No new figures or data.

