## Supplemental Figures for "Spatial transcriptional signatures define margin morphogenesis along the proximal-distal and medio-lateral axes in the complex leaf of tomato (*Solanum lycopersicum*)"

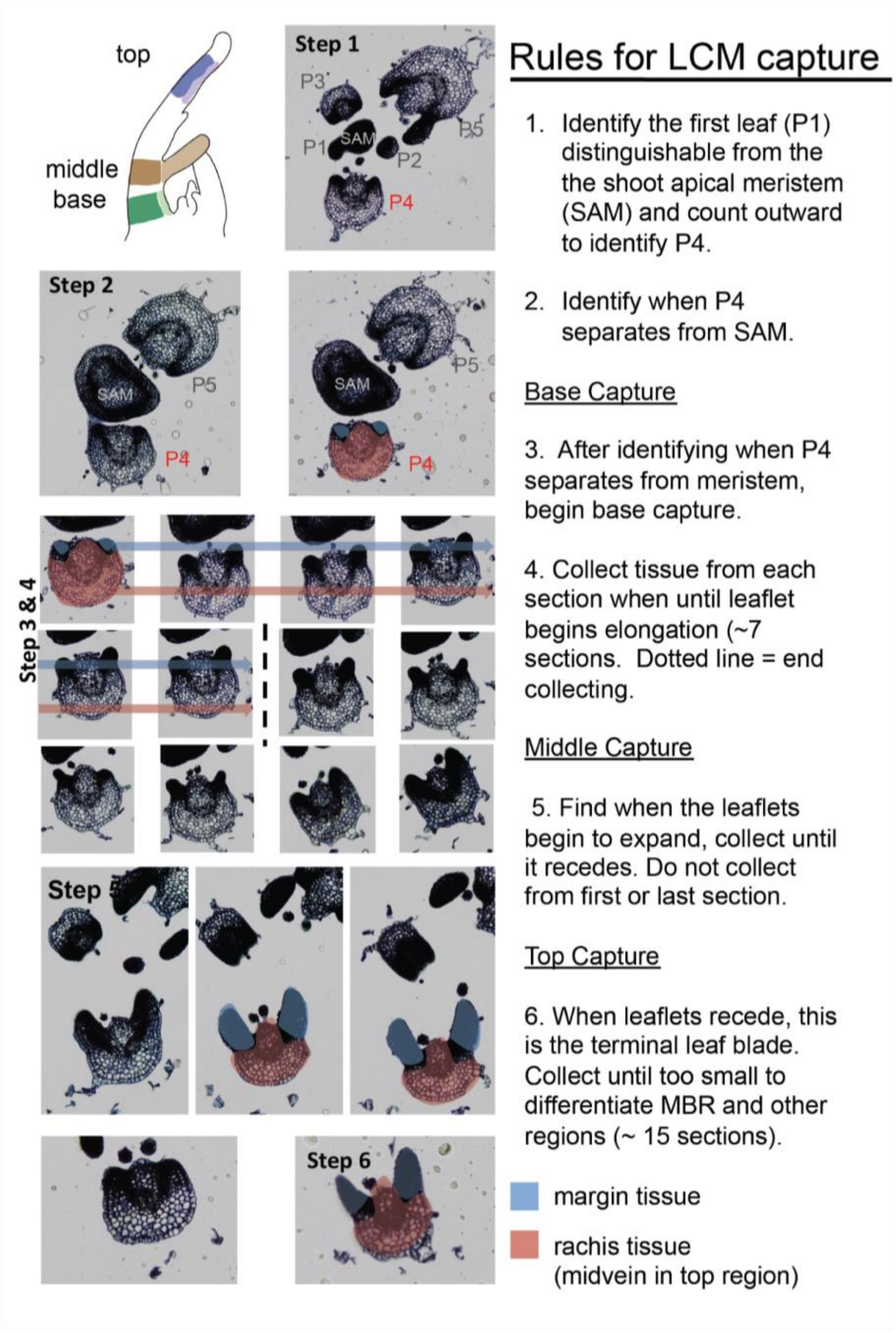


**S1 Figure - Rules for tissue collection via Laser Capture Microdissection (LCM)**
Schematic and rules that were followed when collecting tissues using an LCM microscope. In the top left corner \the colors highlight the separation of the margin (lighter colors) and rachis (darker colors) along the top (purple), middle (brown), and base (green).

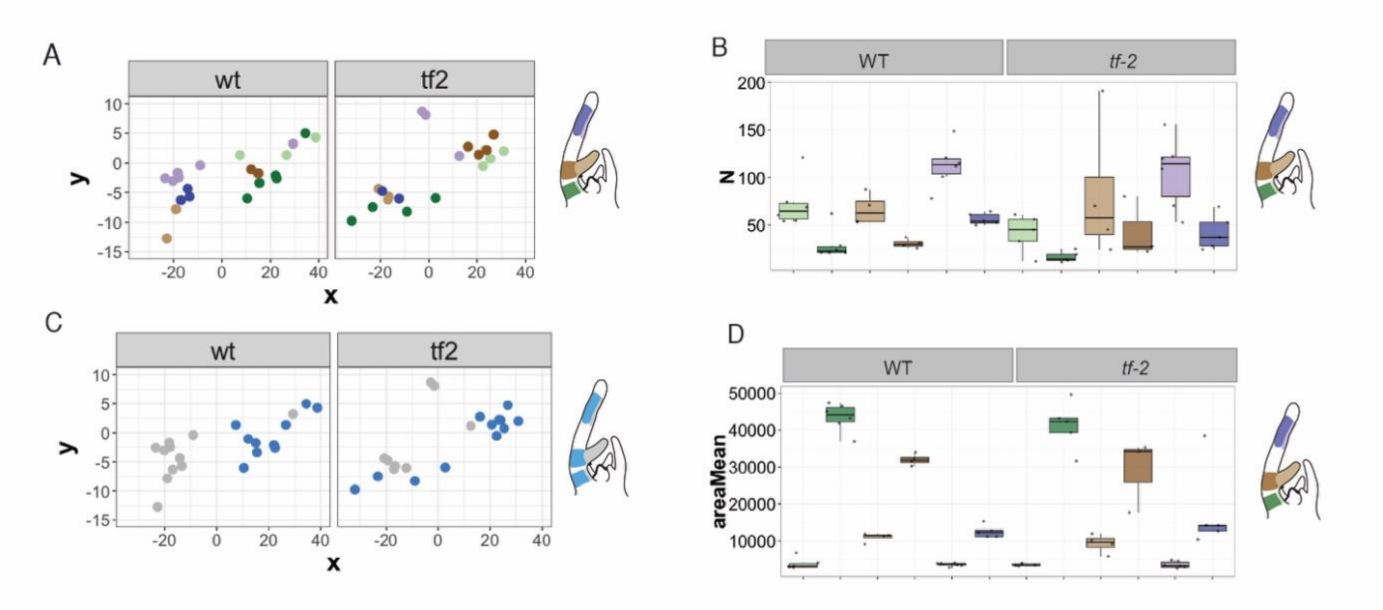


**S2 Figure - Laser cutting to obtain sufficient amounts of RNA for RNA amplification and Illumina sequencing**

Multiple Dimensional Scaling (MDS) analysis of gene expression patterns of normalized reads, colored by (A) sample type and (C) margin (gray) and rachis (blue) tissue.

The type and amount of tissue needed varied depending on cellular density and the amount of tissue of each developmental stage on a P4 leaf primordium. Boxplots displaying the number of LCM (B) cuts and (D) area (µm) needed per tissue sub-region to obtain enough tissue for RNA amplification and Illumina sequencing. (A), (B), and (D), colors highlight the separation of the margin (lighter colors) and rachis (darker colors) along the top (purple), middle (brown), and base (green).

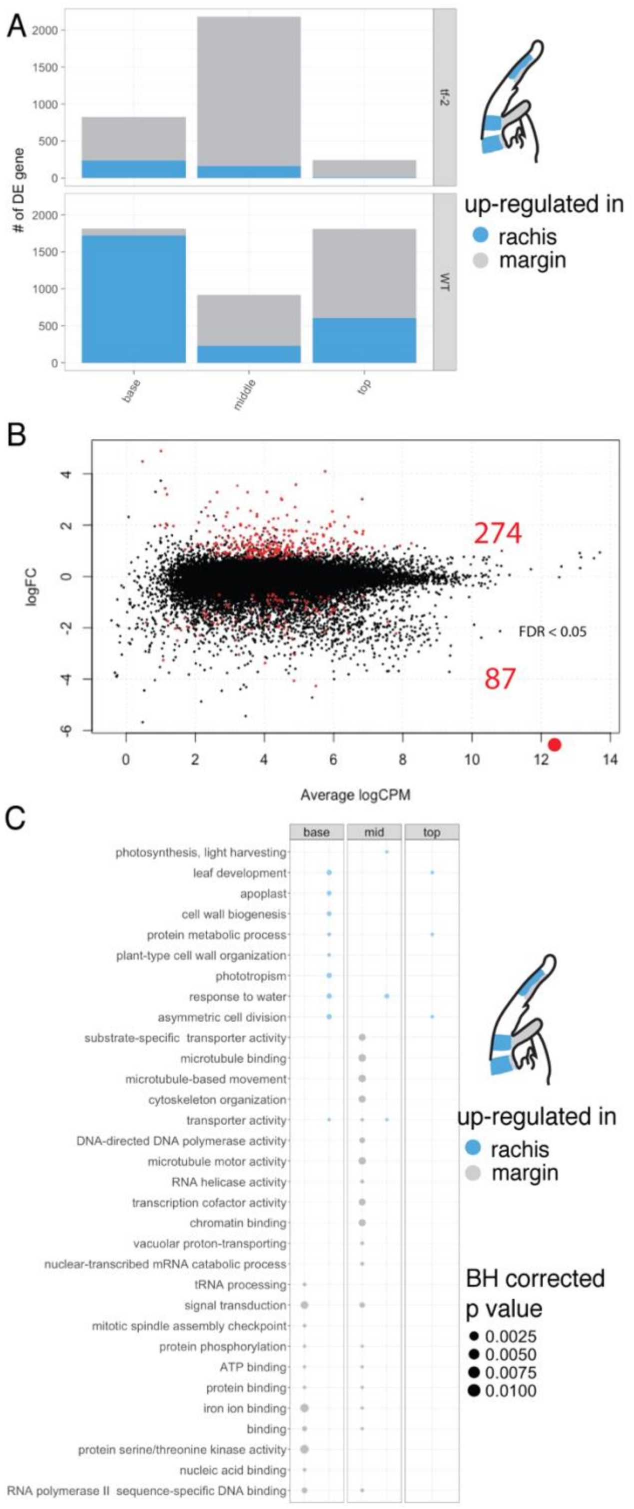


**S3 Figure - Summary of the results of differential gene expression analysis in wild type and *tf-2***

(A) Bar graph displaying the number of differentially expressed genes in wild type and *tf-2* margin (grey) vs. rachis tissue (blue) based on all analyses. Each differential expression analysis was performed separately for each genotype and in each region (top, middle, and base) between the margin and rachis. (B) Results of differential gene expression analysis of *tf-2* displaying average log Counts Per Million (LogCPM) over log Fold Change (logFC), illustrating the number of significant differentially regulated genes (red) between margin and rachis tissue in the top, middle, and base. (C) Pointplot displaying the summary of GO terms describing up-regulated genes in the margin (gray) compared to rachis tissue (blue) in *tf-2*. See **Dataset S10.**

A

| | PC1 | PC2 | PC3 | PC4 | PC5 | PC6 |
|---|---|---|---|---|---|---|
| Standard deviation | 1.3841 | 1.2535 | 1.0666 | 0.9009 | 0.7508 | 1.65E-15 |
| Proportion of Variance | 0.3193 | 0.2619 | 0.1896 | 0.1353 | 0.09395 | 0.00E+00 |
| Cumulative Proportion | 0.3193 | 0.5812 | 0.7708 | 0.9061 | 1 | 1.00E+00 |

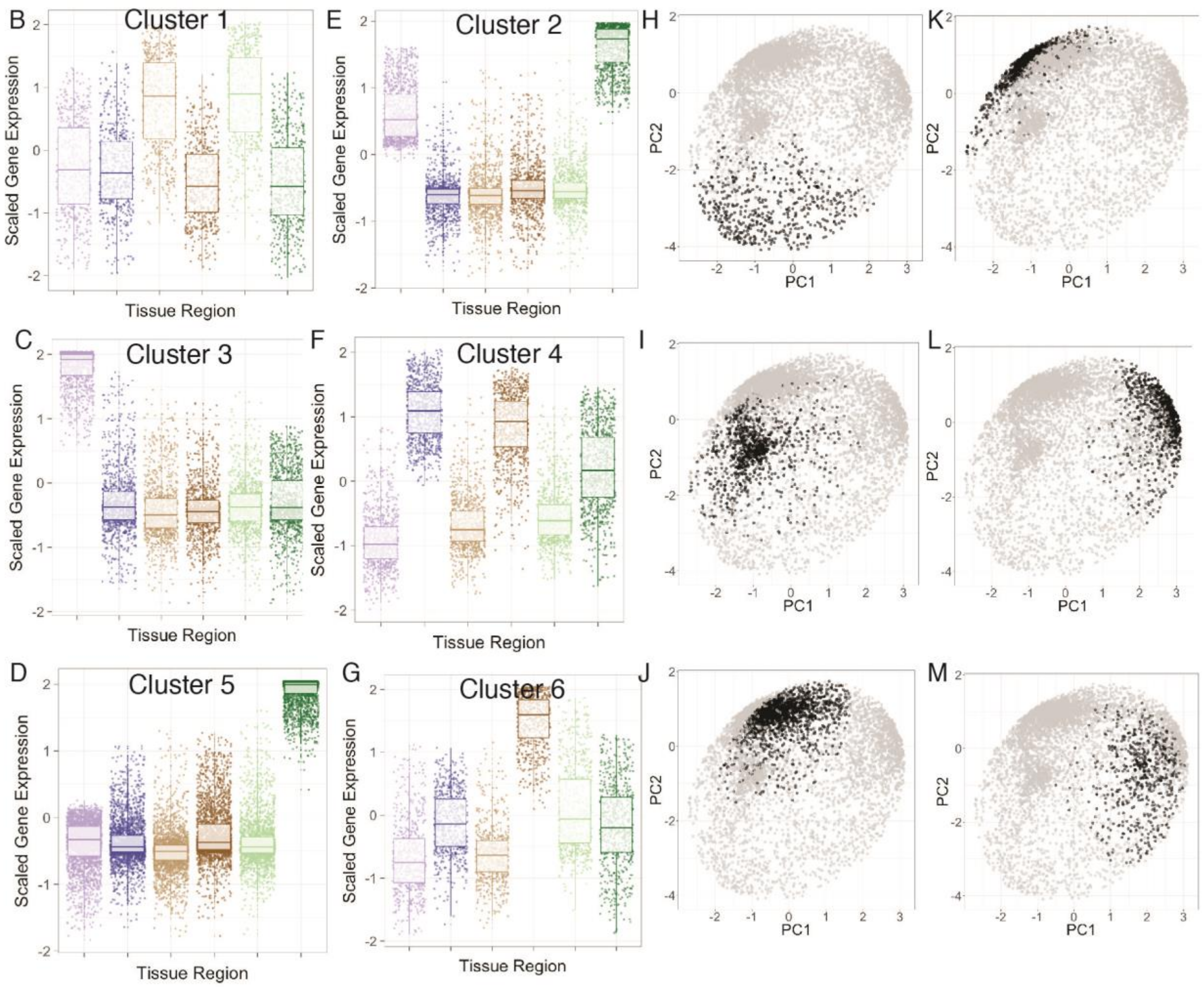


**S4 Figure - Relationship between SOM clustering analysis and PCA analysis performed on wild-type genes across tissues**

PCA was performed on normalized gene expression counts. SOM clustering was performed using the same normalized gene expression counts, with a defined clustering map size of six. (A) Statistics of PCA results. (B) - (G) summarize each cluster, where each point represents a gene in the cluster and its expression value across the six tissue regions. (H) - (M) The expression of each gene in the PC space; bold points reflect the genes present in each cluster based on SOM analysis. (B) - (G) Colors highlight the separation of the margin (lighter colors) and rachis (darker colors) along the top (purple), middle (brown), and base (green).

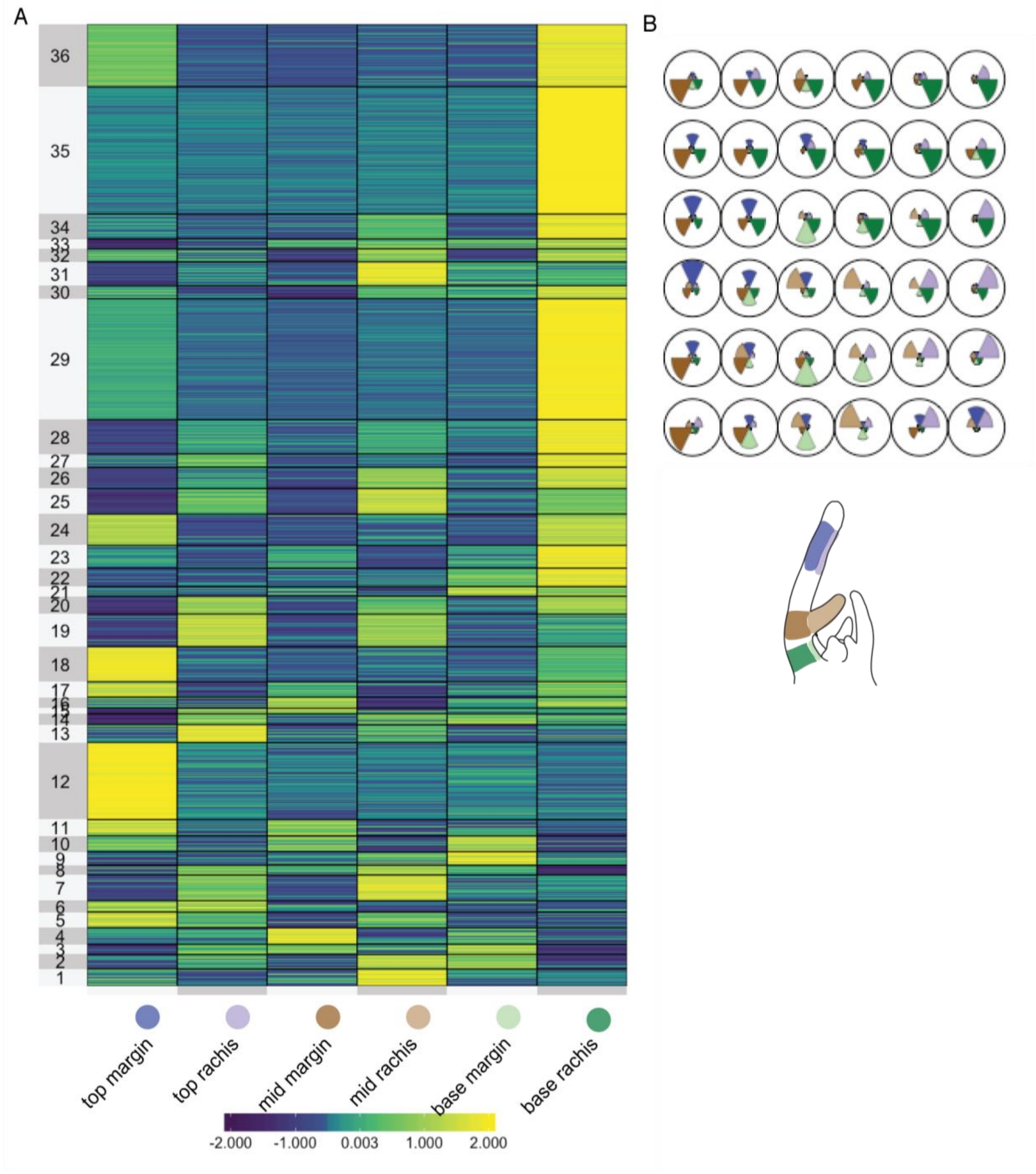


**S5 Figure - Visualization of large SOM clustering analysis**
(A) Heatmap representing the gene expression patterns of the 36 gene clusters. (B) Codevector map displaying the contribution of the expression of each gene in a specific tissue to cluster assignments.

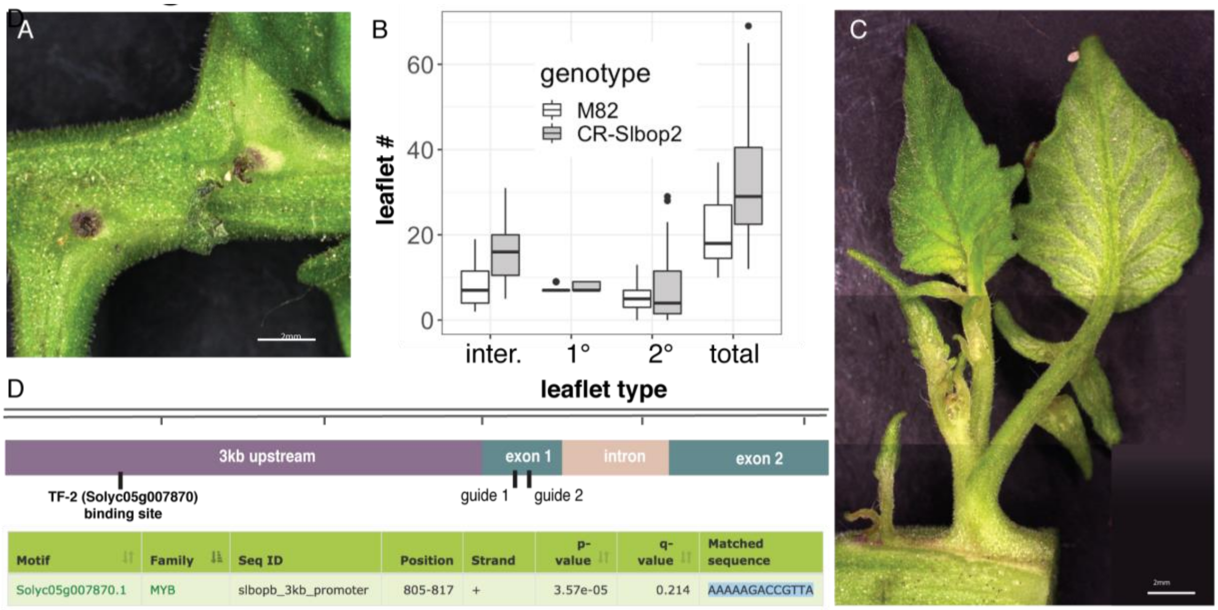


**S6 Figure - Phenotyping of *CR-slbop2* and genomic map of *SlBOP2***

(A) Scars following ectopic shoot apical meristem (SAM) tissue death at the leaflet nodes of a *CR-slbop2* leaf. (B) Leaf complexity counts in wild type (M82) and *CR-slbop2* plants (C) Example of an ectopic SAM that is mature enough to contain complex leaves. (D) *SlBOP2* genomic region showing a TF-2 binding site and the locations of the CRISPR guides used to create the *CR-slbop2* line. Scale bar = 2 mm in (A) and (C).
